# Maternal diet disrupts the placenta-brain axis in a sex-specific manner

**DOI:** 10.1101/2021.11.12.468408

**Authors:** Alexis M Ceasrine, Benjamin A Devlin, Jessica L. Bolton, Lauren A. Green, Young Chan Jo, Carolyn Huynh, Bailey Patrick, Kamryn Washington, Cristina L. Sanchez, Faith Joo, A. Brayan Campos-Salazar, Elana R. Lockshin, Cynthia Kuhn, Susan K. Murphy, Leigh Ann Simmons, Staci D. Bilbo

**Author notes:** Correspondence and requests for materials should be addressed to Staci D. Bilbo.

## Abstract

High maternal weight is associated with a number of detrimental outcomes in offspring, including increased susceptibility to neurological disorders such as anxiety, depression, and communicative disorders (e.g. autism spectrum disorders)^1–8^. Despite widespread acknowledgement of sex-biases in the prevalence, incidence, and age of onset of these disorders, few studies have investigated potential sex-biased mechanisms underlying disorder susceptibility. Here, we use a mouse model to demonstrate how maternal high-fat diet, one contributor to overweight, causes endotoxin accumulation in fetal tissue, and subsequent perinatal inflammation influences sex-specific behavioral outcomes in offspring. In male high-fat diet offspring, increased macrophage toll like receptor 4 signaling results in excess phagocytosis of serotonin neurons in the developing dorsal raphe nucleus, decreasing serotonin bioavailability in the fetal and adult brain. Bulk sequencing from a large cohort of matched first trimester human fetal brain, placenta, and maternal decidua samples reveals sex-specific transcriptome-wide changes in placenta and brain tissue in response to maternal triglyceride accumulation (a proxy for dietary fat content). Further, we find that fetal brain serotonin levels decrease as maternal dietary fat intake increases in males only. These findings uncover a microglia-dependent mechanism through which maternal diet may impact offspring susceptibility for neuropsychiatric disorder development in a sex-specific manner.

## MAIN

In the United States, more than 50% of women are overweight or obese when they become pregnant^9^, and elevated maternal weight is associated with adverse outcomes in offspring, particularly increased incidence of neuropsychiatric disorders such as anxiety, and depression^1–8^. Diets high in saturated fats are one major contributor to overweight and obesity^10^. Thus, we sought to dissect the contribution of a maternal high-fat diet (mHFD) to offspring risk for neuropsychiatric disorder development.

Rodent and non-human primate models have implicated perinatal high-fat diet in the establishment of offspring anxiety^11–13^. Decades of research into anxiety and depression have strongly linked disruptions in the serotonin (5-HT) system as causal to neuropsychiatric diseases such as anxiety and depression. Although women disproportionately bear the burden of these disorders^14, 15^, little has been done to investigate the contribution of 5-HT to anxiety/depression on the basis of sex. Understanding the developmental mechanisms through which sex contributes to neuropsychiatric disorder development, susceptibility, and severity in response to environmental exposures such as diet is of utmost importance when considering clinical implications such as treatment efficacy.

### Maternal high-fat diet imparts sex-specific offspring behavioral outcomes

To interrogate the contribution of maternal dietary fat content to offspring neurological development, female wild-type (*C57BL/6J*) mice were placed on either a high-fat (45% kcal from fat; HFD) or low-fat (10% kcal from fat, sucrose matched; LFD) diet at approximately 4 weeks of age (juvenile) for six weeks prior to mating (Fig 1a). Starting females on HFD during the juvenile period best mimics trends observed in fat consumption in human populations – juveniles tend to get more calories from high-fat foods, and their eating patterns as adults tend to mimic their adolescent eating patterns^16–20^. HFD females gained significantly more weight than LFD females before pregnancy and their weight remained elevated throughout gestation (Extended Data Fig 1a-b). No significant differences were observed in litter size, composition, or maternal care (during either the light or dark phase; Extended Data Fig 1c-e). Both male and female maternal high-fat diet (mHFD) offspring weighed significantly more than maternal low-fat diet (mLFD) offspring throughout life, though placental weight at embryonic day 14.5 (e14.5) was not affected by mHFD (Extended Data Fig 1f-g). Offspring behavior was assessed at neonatal, juvenile, and adult time-points (Fig 1a).

**Fig 1.**
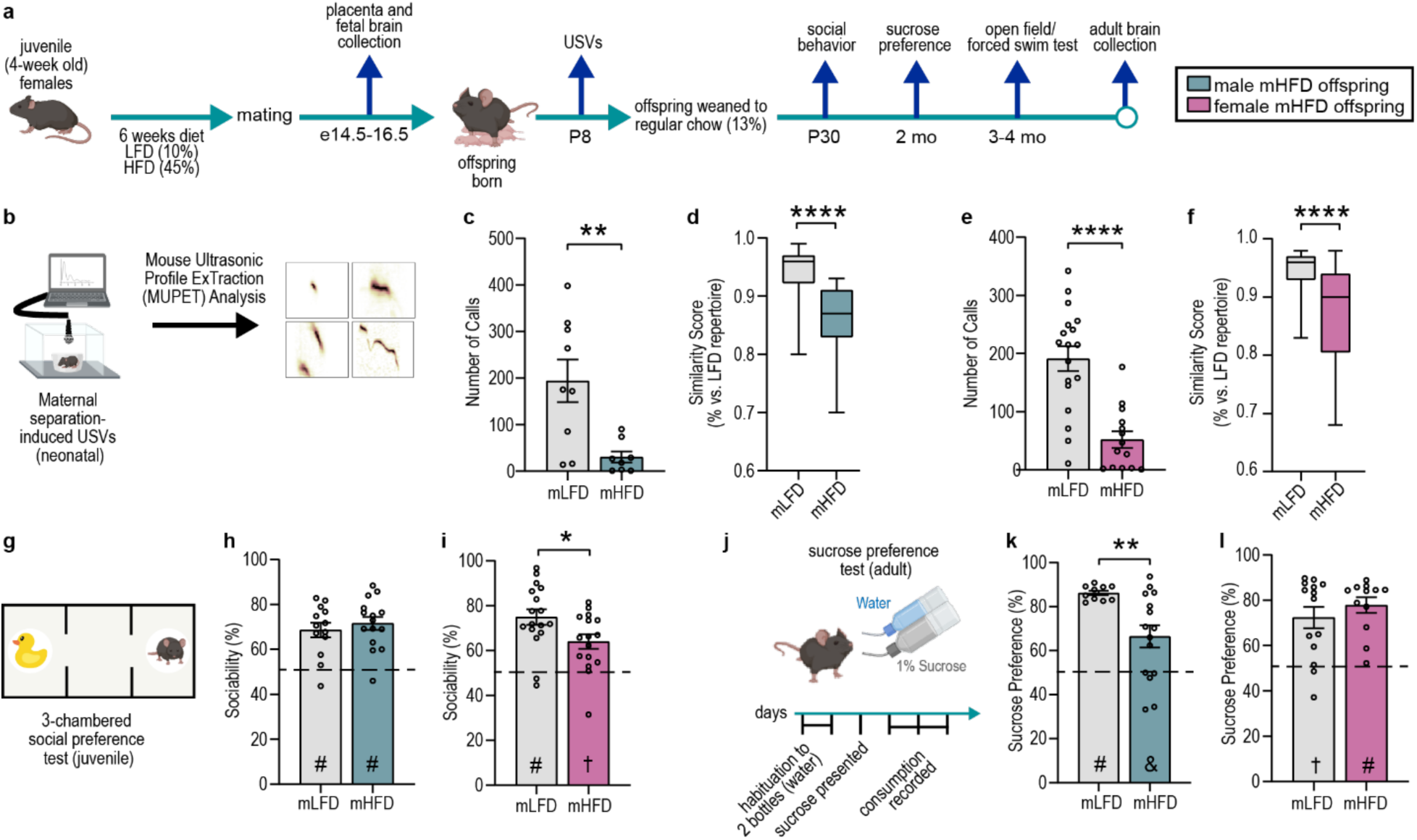
Maternal high-fat diet imparts sex-specific offspring behavioral outcomes. **a**, Schematic of maternal low-fat and high-fat diet paradigm (10% and 45% represent %kcal from fat In diet), **b,** Schematic of neonatal maternal separation-induced ultrasonic vocalization (USV) recording and analysis, **c-f**, mHFD decreases neonatal USV number and syllable repertoire similarity In male and female offspring (n=9 mLFD, 8 mHFD male offspring; 18 mLFD, 14 mHFD female offspring from 5 mLFD and 4 mHFD litters), **g**, Schematic of 3-chambered social preference test, **h,** Male mLFD and mHFD offspring display a strong social preference (n=13 mLFD, 15 mHFD male offspring from 5 mLFD and 6 mHFD litters), **i,** Female mHFD offspring have a reduced social preference versus mLFD offspring (n=17 mLFD, 16 mHFD female offspring from 6 mLFD and 6 mHFD litters), **j**, Schematic of sucrose preference test, **k**, Male mHFD offspring display a decreased preference for sucrose versus mLFD offspring (π=10 mLFD, 15 mHFD male offspring from 5 mLFD and 5 mHFD litters). **I**, Female mLFD and mHFD offspring display a strong sucrose preference (n=14 mLFD, 12 mHFD female offspring from 5 mLFD and 5 mHFD litters). Data are mean ± s.e.m.; asterisk denoted p-values are derived from unpaired two-tailed t-tests (**c, e, i, k**) or paired two-tailed t-tests (**d, f**). other p-values (# p<0.0001, f p<0.001, & p<0.01) are derived from one-sample t-tests assessing difference from chance (50%) **(h, I, k, I).**

Neonatal maternal separation-induced ultrasonic vocalizations (USVs) provide a readout of early communicative behaviors in mice, and peak at postnatal day 8 in *C57BL/6J* mice^21, 22^. USV analyses (Fig 1b) revealed a significant decrease in the number of USVs emitted, total call time, mean call length, and mean syllable number along with a significant increase in inter-syllable interval in both male (Fig 1c, Extended Data Fig 2a-e) and female (Fig 1e, Extended Data Fig 2g-k) mHFD offspring compared to same sex mLFD offspring. Female, but not male, mHFD offspring USVs had a higher mean frequency (kHz) compared to mLFD offspring (Extended Data Fig 2d, j). To rule out a potential shift in USV peak in mHFD offspring, we quantified USVs at P7 and P10 in addition to P8, but mHFD offspring called less across the assessed neonatal window (Extended Data Fig 2f, l). In-depth characterization of overall USV composition (integrating alterations in frequency (kHz), energy (dB), and syllable complexity [e.g. number of syllables, frequency of calls]) using Mouse Ultrasonic Profile ExTraction (MUPET^23^) revealed a significant decrease in USV similarity (an integrated measure of call parameters, see methods for more detail) between mLFD and mHFD male (Fig 1d) and female (Fig 1f) offspring. While less is known about what USV complexity may represent in neonates, adult USVs are context-dependent, and specific USV parameters such as frequency (kHz) are known to vary in response to a stressful environment^24^. To parse out whether the USV changes were reflective of an overall lack of social engagement in mHFD offspring, we assessed juvenile sociability using a 3-chamber social preference test (Fig 1g). Male mHFD offspring showed no change in sociability, time investigating the social stimulus, or time in the social chamber when compared to controls (Fig 1h; Extended Data Fig 2m). Female mHFD offspring had a significantly decreased sociability score, driven by female mHFD offspring spending significantly less time investigating the social stimulus as compared to controls (Fig 1i; Extended Data Fig 2o). Interestingly, USV emission (number of calls) at P8 significantly positively correlated with social behavior in females only (r^2^=0.39, p=0.006 for females, r^2^=0.004, p=0.78 for males), suggesting that decreased USVs may predict impaired social behavior in females, but not in males. In a complementary social novelty preference (SNP) assay, male mHFD showed no change in SNP, time investigating the novel social stimulus, or time in the novel social chamber compared to mLFD controls (Extended Data Fig 2n). Female mHFD offspring had a significantly decreased SNP score, driven by female mHFD offspring spending significantly less time investigating the novel social stimulus and less time in the novel social stimulus chamber compared to mLFD controls (Extended Data Fig 2p). To better understand the behavioral changes affecting male mHFD offspring, we determined average sucrose preference over a 3-day free-choice test (Fig 1j). Mice normally display a strong preference for sucrose, and diminished sucrose preference is one measure of anhedonia, or lack of reward/pleasure. Average sucrose preference was significantly decreased in male, but not female, mHFD offspring (Fig 1k-l). Neither male nor female mHFD offspring showed a change in total liquid consumption (water + sucrose) over the 3-day testing period (Extended Data Fig 3a,f), and there was no correlation between body weight and sucrose preference within either mLFD or mHFD groups. We additionally assessed mobility behavior in a forced swim test (FST; Extended Data Fig 3b). Male mHFD offspring spent significantly less time immobile, moved significantly further, and with a higher velocity than mLFD offspring (Extended Data Fig 3c-e). No changes were observed in FST behavior in female mHFD offspring compared to female mLFD offspring (Extended Data Fig 3g-i). Lastly, neither male nor female mHFD offspring had any changes in general activity or anxiety-like behavior in an open field test (OFT; Extended Data Fig 3j-r). This suggests that the social deficits observed in females are not due to impaired motor skills or an anxiety-like phenotype, and that the increased velocity and distance moved by males in the FST are not due to general hyperactivity. In sum, despite some similar behavioral phenotypes in USVs in early life, by adolescence and into adulthood the behavior patterns diverge based on sex, with mHFD female offspring showing social deficits, and mHFD male offspring showing diminished non-social reward behavior and increased activity in the FST.

### mHFD decreases 5-HT in male offspring and increasing 5-HT levels is sufficient to prevent behavioral phenotypes

We observed altered neonatal USVs, anhedonia, and increased activity during the FST in male mHFD offspring. These behaviors are associated with diminished brain 5-HT (5-HT) in mice^25–29^. Previous research has demonstrated that brain 5-HT levels during discrete windows of development are dependent on placental 5-HT synthesis^30^. The placenta is a temporary organ that develops during early gestation and surrounds the fetus to both restrict and facilitate macromolecule and nutrient transport. Further, the placenta acts as a critical interface between the maternal environment and the developing fetal brain and responds to maternal diet/environment in a fetal sex-specific manner^31–33^. However, a potential role for fetal sex in placental 5-HT production is unknown. Assessment of 5-HT levels in mLFD and mHFD placenta and fetal forebrain revealed a male-specific decrease in tissue 5-HT in mHFD offspring (Fig 2a-d, also see Extended Data Fig 4d-g). Circulating maternal 5-HT levels were not changed by HFD, suggesting that this regulation is within fetal tissue (Extended Data Fig 4a). In agreement with previous studies demonstrating that placental 5-HT correlates with fetal brain 5-HT^30^, we observed a significant positive correlation between placental and fetal forebrain 5-HT in mLFD offspring (Fig 2e, see also Extended Data Fig 5f). Interestingly, the correlation was exclusively driven by male 5-HT levels, but mHFD disrupted the correlation between placental and brain 5-HT levels in males, potentially a floor effect due to low 5-HT levels (Fig 2f, see also Extended Data Fig 5g).

**Fig 2.**
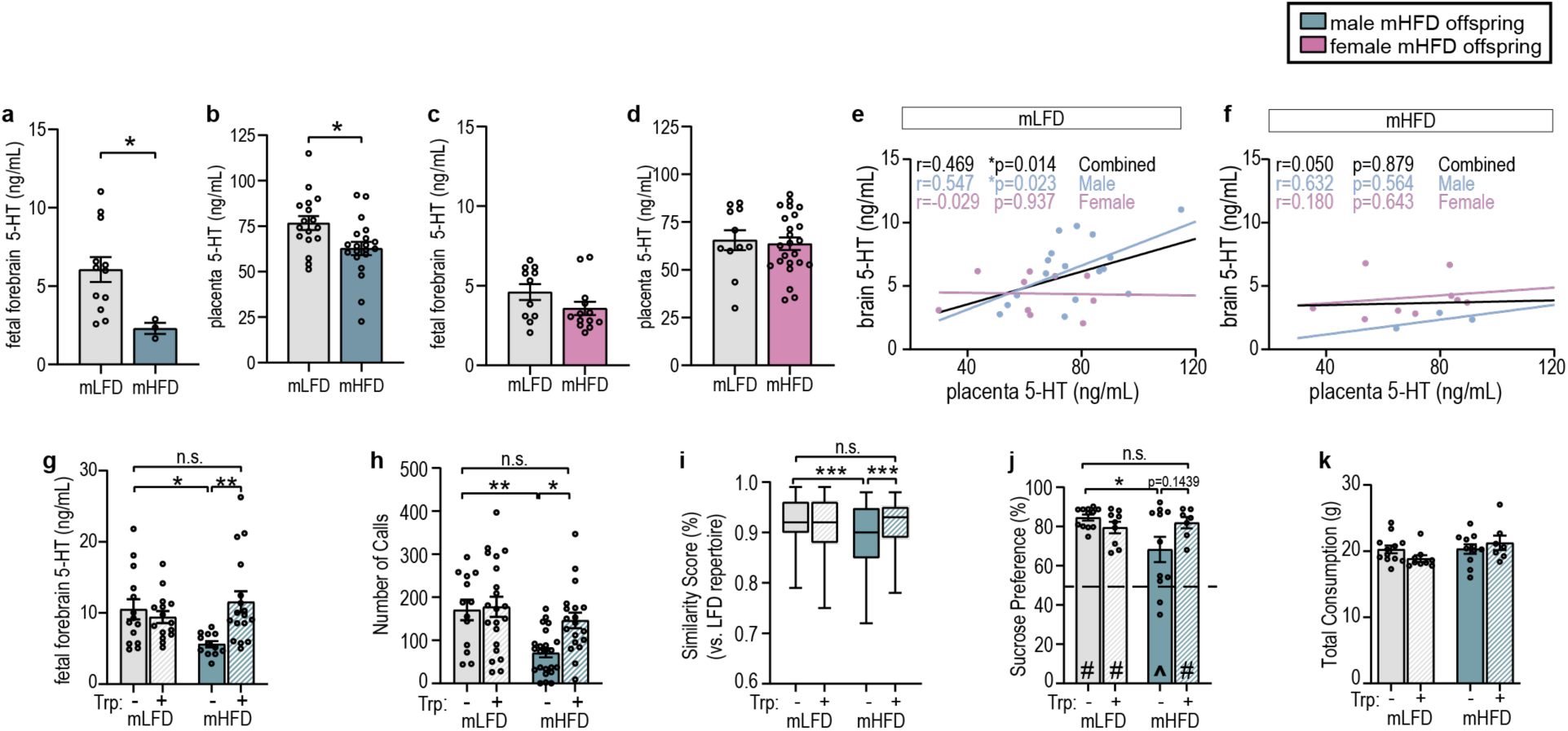
mHFD decreases 5-HT in male offspring and increasing 5-HT level is sufficient to prevent behavioral phenotypes. **a**, Male mHFD offspring have decreased fetal forebraln serotonin (n=12 mLFD, 3 mHFD male offspring from 3 mLFD and 3 mHFD litters), **b**, Male mHFD offspring have decreased placenta serotonin (n=18 mLFD, 20 mHFD offspring from 4 mLFD and 6 mHFD litters), **c-d**, mHFD does not Impact fetal serotonin levels In female offspring (forebraln: n=11 mLFD, 13 mHFD from 3 mLFD and 3 mHFD litters; placenta: n=11 mLFD, 24 mHFD female offspring from 4 mLFD and 6 mHFD litters) **e-f**, Fetal serotonin levels are significantly correlated between brain and placenta In mLFD males only (n=17 male, 10 female mLFD offspring from 4 mLFD litters; 3 male, 9 female mHFD offspring from 3 mHFD litters), **g,** Maternal dietary tryptophan enrichment Increases serotonin levels In mHFD fetal male forebraln (n=14 mLFD, 15 mLFD+Trp, 12 mHFD, 17 mHFD+Trp offspring from 5 mLFD, 5 mLFD+Trp, 3 mHFD, and 5 mHFD+Trp litters) **h-i**, Maternal dietary tryptophan enrichment rescues mHFD-lnduced decrease In USV number and syllable repertoire similarity (n= 13 mLFD, 21 mLFD+Trp, 23 mHFD, 19 mHFD+Trp offspring from 5 mLFD, 7 mLFD+Trp, 8 mHFD, and 7 mHFD+Trp litters), **j-k**, Maternal dietary tryptophan enrichment rescues sucrose preference In mHFD male offspring and does not affect total consumption (n= 12 mLFD, 9 mLFD+Trp, 11 mHFD, 7 mHFD+Trp male offspring from 6 mLFD, 5 mLFD+Trp, 5 mHFD, and 6 mHFD+Trp litters). Data are mean ± s.e.m. asterisk denoted p-values are derived from unpaired two-tailed t-tests (**a, b**), Pearson’s correlation (**e, f**), or two-way ANOVA (fat content x tryptophan content; **g, h, i, j**). other p-values (# p<0.0001, ^p<0.05) are derived from one-sample t-tests assessing difference from chance (50%) (j). n.s. not significant.

Quantitative real-time PCR (qPCR) showed a decrease in both the 5-HT synthesis enzyme tryptophan hydroxylase 2 (*Tph2*) and the 5-HT transporter (*5HTT*) in male mHFD placenta (Extended Data Fig 4b-c). To better understand the dynamics of placental 5-HT in response to mHFD, we additionally quantified levels of 5-HT, the 5-HT precursor tryptophan, and the 5-HT metabolite 5-hydroxyindoleacetic acid (5-HIAA), in the placenta using high-performance liquid chromatography (HPLC; Extended Data Fig 4d-g and ExtendedDataFig4_RawData_Stats). In agreement with decreased *Tph2* levels, 5-HT synthesis (ratio of 5-HT to tryptophan) was significantly decreased in male, but not female, mHFD offspring placenta (Extended Data Fig 4d, f). We did not observe any changes in the expression of the alternate 5-HT synthesis enzyme tryptophan hydroxylase 1 (*Tph1*), monoamine oxidase A (*MAOA*), an enzyme responsible for the oxidative deamination of neurotransmitters, including 5-HT, or the rate-limiting enzyme indoleamine 2,3-dioxygenase (*Ido1*) that promotes the degradation of tryptophan through the kynurenine pathway in male placentas (Extended Data Fig 4b). In line with this, we also did not observe any changes in the protein levels of quinolinic acid, and kynurenic acid levels were undetectable in e14.5 brain and placental tissue, suggesting that tryptophan is not being diverted into the kynurenine pathway in mHFD offspring (Extended Data Fig4h-i). Comparing the ratio of 5-HIAA to 5-HT (5-HT turnover) revealed a modest but significant increase in 5-HT turnover in male mHFD offspring placenta, suggesting that in addition to diminished 5-HT synthesis, increasing turnover may contribute to total decreased placental 5-HT levels in male mHFD offspring (Extended Data Fig 4d). In the fetal forebrain, we did not see changes in 5-HT synthesis or turnover, but we did see a decrease in tryptophan in both male and female mHFD offspring (Extended Data Fig 4e, g). Lastly, *MAOA* expression was significantly decreased in female mHFD placenta, suggesting that an alternate neurotransmitter may be affected in female mHFD offspring, and that the placenta may play a critical role as a source for brain neurotransmitters other than 5-HT (Extended Data Fig 4c).

To determine if increasing 5-HT in male mHFD offspring could rescue the mHFD-induced behavioral outcomes, we sought to rescue 5-HT by supplementing the maternal low- and high-fat diets with tryptophan. Tryptophan is the precursor to 5-HT, and placental 5-HT levels are largely dependent on maternal tryptophan levels^30, 32^. Tryptophan supplementation (+Trp; provided in LFD or HFD chow for the same duration as in Fig 1a) did not affect maternal weight gain, litter composition (i.e., size and sex ratio), or offspring weights (Extended Data Fig 5a-d).

Maternal tryptophan supplementation increased placental 5-HT levels in male mLFD offspring (Extended Data Fig 5e) in support of previous literature demonstrating that placental 5-HT levels are dependent on maternal tryptophan levels. In male mHFD offspring, tryptophan supplementation was unable to substantially increase placental 5-HT levels (Extended Data Fig 5e). Conversely, maternal tryptophan supplementation did increase fetal forebrain 5-HT levels in male mHFD, but not mLFD offspring (Fig 2g). Further, Pearson correlation analyses demonstrated that maternal tryptophan enrichment disrupted the significant positive correlation between brain and placental 5-HT levels in male mLFD offspring (Extended Data Fig 5f). No significant correlation was detected in mHFD (in agreement with our previous finding) or mHFD+Trp offspring (Extended Data Fig 5g). This suggests that in mHFD+Trp offspring, the fetal brain produces 5-HT independently of the placenta. Together these data demonstrate that in males, placental 5-HT levels are dependent on maternal tryptophan levels and can be disrupted by maternal diet, but fetal forebrain 5-HT levels are not solely dependent on placental 5-HT levels. Further, mHFD led to decreased 5-HT levels in adult male midbrain – where the majority of 5-HT neuronal cell bodies reside – levels that were also rescued by maternal tryptophan enrichment (Extended Data Fig 5h).

To determine if mHFD-induced behavioral deficits in male offspring were rescued by maternal tryptophan supplementation, we assessed USVs and sucrose preference in male mLFD and mHFD offspring. Tryptophan supplementation did not impact USV or sucrose preference behavior in male mLFD offspring (Fig 2h-k, Extended Data Fig5 i-m). However, in mHFD offspring, maternal tryptophan supplementation fully rescued the decreased USV call number and similarity, as well as the increased inter-syllable interval (Fig 2h-i, Extended Data Fig 5k), partially rescued the deficits seen in total USV call time, mean call length, and mean syllable number (Extended Data Fig5 i-m), as well as the diminished sucrose preference (Fig 2j-k). In all cases, mHFD+Trp offspring behavior was indistinguishable from that of mLFD or mLFD+Trp offspring. Of note, maternal tryptophan supplementation rescued male mHFD offspring USV phenotypes despite the observation that male mHFD+Trp offspring still weigh more than mLFD and mLFD+Trp offspring (see Extended Data Fig 5c), suggesting that changes in offspring weight is not driving the USV phenotypes. Further, maternal tryptophan supplementation did not alter male offspring social behavior or open field behavior (Extended Data Fig 5n-u).

Although we did not expect to see a rescue of behavioral phenotypes in female mHFD+Trp offspring because mHFD did not impact 5-HT levels in females, we assessed midbrain 5-HT and behavior outcomes. Adult female offspring midbrain 5-HT levels were unaffected by mHFD or tryptophan supplementation (Extended Data Fig 6a). Maternal tryptophan enrichment did not rescue female mHFD offspring USV behavior, but interestingly did decrease USV call similarity and call length in mLFD offspring (Extended Data Fig 6b-h). Maternal tryptophan supplementation also increased USV frequency (kHz) in female offspring regardless of maternal dietary fat content (Extended Data Fig 6g). This suggests that despite similar neonatal behavioral outcomes in male and female mHFD offspring, the mechanisms driving USV behavior in males and females may be distinct. Interestingly, maternal tryptophan enrichment increased sucrose preference in female offspring regardless of maternal dietary fat intake (Extended Data Fig 6i). Maternal tryptophan enrichment did not rescue female mHFD social behavior deficits (Extended Data Fig 6j-m). In sum, these data demonstrate 1) a developmental role for 5-HT in males that is perturbed by maternal high-fat diet, and 2) that maternal tryptophan enrichment is sufficient to rescue mHFD-induced brain 5-HT deficiency and associated 5-HT-dependent behaviors in males.

### mHFD induces Tlr4-dependent inflammation driving offspring behavior changes

Maternal HFD creates an environment of chronic inflammation that affects the developing fetus as well as the placenta^34^. Growing literature has demonstrated complex relationships between inflammation and 5-HT^35–37^, and we hypothesized that the mHFD-induced inflammatory environment is translated to the fetus, and responsible for decreased 5-HT in male mHFD offspring. At e14.5, we observed increased macrophage (F4/80+ cells located on the fetal side of the placenta) density in male and female mHFD placenta (Extended Data Fig 7a), as well as increased Iba1 and CD68 immunoreactivity (labeling microglia and phagosomes, respectively) in both male and female fetal dorsal raphe nuclei (DRN), where the majority of serotonergic cell bodies reside (Fig 3a). Microglia influence central circuits in numerous ways, such as by pruning excess synapses or phagocytosing excess or dying progenitor cells^38–41^. Interestingly, qPCR for genes important for phagocytosis from isolated midbrain microglia at e14.5 did not reveal any significant changes in gene expression, although the levels of some genes were near or below the limit of detection (see Methods, Extended Data Fig 7b). Given the increased immunoreactivity seen by immunohistochemistry (IHC), we further investigated potential interactions between microglia and the developing central 5-HT system by 3D reconstructing microglia from e14.5 DRN. IMARIS reconstructions revealed that 5-HT signal comprised a higher percentage of phagosomes (CD68) in male mHFD microglia *versus* mLFD (Fig 3b), demonstrating that microglial phagocytosis of serotonergic neurons is increased in males in the context of mHFD, leading to decreased central 5-HT bioavailability. Postnatal microglia express the 5-HT receptor 5HT2B^42^ and decrease their phagocytic capacity in response to 5-HT^43^. Further, mice lacking microglial 5HT2B display prolonged neuroinflammation in response to lipopolysaccharide (LPS), a classic agonist of macrophage toll-like receptor signaling^44^. Thus, we sought to investigate potential feedback between 5-HT levels and microglial phagocytosis of 5-HT neurons. Microglial 5-HT phagocytosis was rescued in mHFD+Trp offspring, in parallel with 5-HT levels (Fig 3b; see Figure 2g), suggesting that in the context of mHFD, enrichment of 5-HT levels is sufficient to prevent further aberrant microglia phagocytosis. Consistent with our observations that mHFD does not decrease 5-HT levels in females, we did not see increased 5-HT signal within phagosomes in mHFD female microglia (Extended Data Fig 7c). Interestingly, there was a significant increase in 5-HT signal within phagosomes in reconstructed microglia from mHFD+Trp female microglia (Extended Data Fig 7c), possibly contributing to the trend towards lower midbrain 5-HT we observed in the adult female mHFD+Trp offspring (see Extended Data Fig 6a). This trend parallels the open field behavior, with mHFD+Trp female offspring showing increased activity in the open field test (see Extended Data Fig 6n-o), suggesting that fetal microglial interactions with 5-HT neurons may dictate different behaviors in males and females.

**Fig 3.**
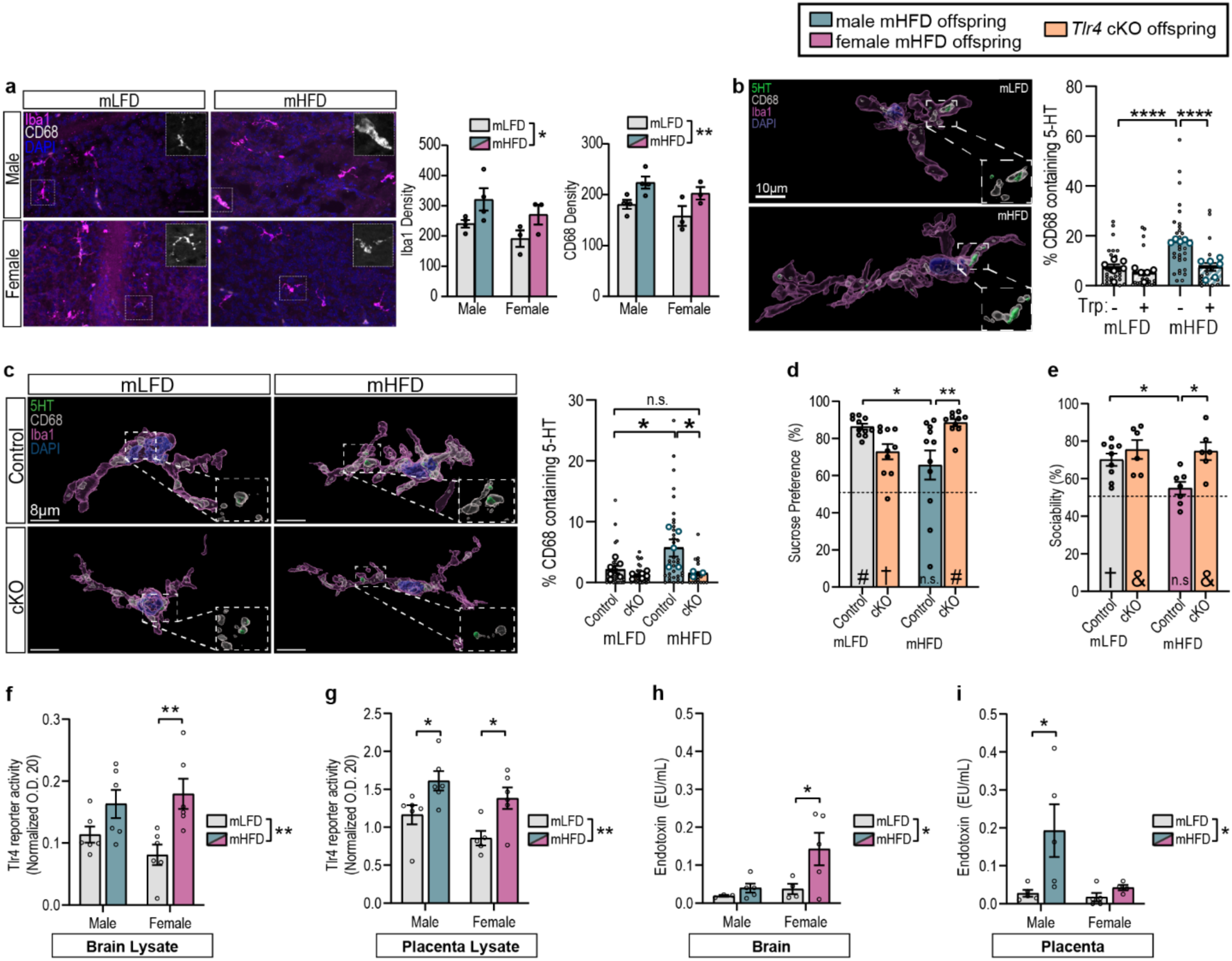
mHFD induces Tlr4-dependent inflammation driving offspring behavior changes. **a**, Increased microglia and phagosome immunoreactivity in male and female mHFD DRN at e14.5 (scale=50pm; n=4 male and 3 female/diet from 3mLFD and 3mHFD litters), **b**, IMARIS reconstructions of DRN microglia reveals Increased microglial phagocytosis of serotonin In males at e14.5. Statistics are displayed for animal average (large circles), small circles are individual microglia reconstructions (n=6 n∩LFD, 5mLFD+Trp, 5 mHFD, 6 mHFD+Trp mice from 3 mLFD, 2 mLFD+Trp, 3 mHFD, and 3mHFD+Trp litters), **c**, IMARIS reconstruction of DRN microglia In male ∞ntrol and *Tlr4* cKO e14.5 offspring (n=6 mLFD control, 6 mLFD cKO, 5 mHFD control, and 3 mHFD cKO male offspring from 7 mLFD and 5 mHFD litters), **d,** Decreased sucrose preference Is rescued In male mHFD offspring lacking macrophage-Tlr4 signaling, (n=11 mLFD, 10 mLFD cKO, 11 mHFD, and 10 mHFD cKOfrom 5 mLFD and 6 mHFD litters), **e**, Decreased social preference Is rescued In female mHFD offspring lacking macrophage-Tlr4 signaling, (n=9 mLFD, 6 mLFD cKO, 7 mHFD, and 6 mHFD cKO from 5 mLFD and 5 mHFD litters), **f-g,** Tlr4-reporter activity In response to e14.5 brain (f) and placenta (g) lysate from mLFD and mHFD offspring (n=6 offsprlng/sex/dlet from 5 mLFD and 5 mHFD litters), **h-i,** Endotoxin levels In mLFD and mHFD brain (h) and placenta (I) tissue lysate (n=3-5 mLFD male, 4-5 mLFD female, 5 mHFD male, 4-5 mHFD female from 5 mLFD and 5 mHFD litters). Data are mean ± s.e.m.; asterisk denoted p-values are derived from two-way ANOVA (**a-l**). other p-values (# p<0.0001, † p<0.001, & p<0.01) are derived from one-sample t-tests assessing difference from chance (50%) (**d, e**). n.s. not significant

The presence of 5-HT receptors on placental macrophages, or the potential influence of 5-HT on placental macrophages is unknown, although recent studies have demonstrated the presence of functional 5-HT transporters in placental tissue^45, 46^. Thus, we sought to investigate crosstalk between placental 5-HT and placental macrophage density as well as fetal brain 5-HT and microglial activity. Maternal tryptophan supplementation did not significantly influence placental macrophage density in either mLFD or mHFD male or female offspring (Extended Data Fig 7d). Considering our finding that male mLFD+Trp offspring have increased placental 5-HT (see Extended Data Fig 5e), this suggests that placental macrophage density is not significantly influenced by 5-HT.

To further investigate the potential pro-inflammatory pathways leading to increased placental macrophage and fetal brain microglia density, we assessed a panel of pro-inflammatory markers with qPCR (Extended Data Fig 7e). This revealed a specific increase in Toll-like receptor 4 (Tlr4) in both male and female mHFD placenta (Extended Data Fig 7e). Tlr4 is a pattern recognition receptor known to initiate pro-inflammatory signaling cascades in response to infectious pathogens (e.g. bacterial, viral), and whose expression is increased in response to saturated fats/obesity^47–50^. Isolated fetal microglia and placental macrophages from obese dams have exaggerated cytokine production in response to lipopolysaccharide (LPS), a classic Tlr4 agonist^33^.

To determine if increased *Tlr4* signaling in macrophages was causal to decreased 5-HT and 5-HT-dependent behaviors in males, or to behavioral changes in females, we generated a conditional Tlr4 knock-out mouse (cKO) line using Cx3cr1-Cre BAC-transgenic^51^ and Tlr4^f/f^ mice^52^ (Extended Data Fig 7f). Female control (*Tlr4^f/f^*) mice were transitioned to HFD for six weeks, then subsequently mated to male *Cx3cr1-Cre;Tlr4^f/f^* mice to generate control and cKO mHFD offspring. qPCR revealed significant (∼95%) knockdown of *Tlr4* in cKO microglia (Extended Data Fig 7f). 5-HT was increased in male fetal forebrain and adult midbrain in mHFD cKO offspring (Extended Data Fig 7g-i). IMARIS reconstructions of mLFD and mHFD microglia from control and *Tlr4* cKO e14.5 DRN revealed a significant rescue of 5-HT phagocytosis by microglia in mHFD *Tlr4* cKO male offspring (Fig 3c). No significant changes were detected in females (Extended Data Fig 7j). Loss of *Tlr4* decreased USV similarity in male mLFD offspring while increasing it in male mHFD offspring (Extended Data Fig 8a), suggesting a role for macrophage *Tlr4* signaling in USV behavior independent of maternal dietary fat composition. In male mHFD offspring, loss of *Tlr4* completely rescued mHFD-induced anhedonia (Fig 3d, Extended Data Fig 8c), and did not influence male social preference (Extended Data Fig 8d). Interestingly, female mHFD *Tlr4* cKO offspring did not display a preference for sucrose (Extended Data Fig 8e), suggesting that increased Tlr4 signaling in female mHFD offspring may be preventing anhedonia. *Tlr4* knockdown partially rescued the number of USVs, but did not rescue total call time, or mean call length in male mHFD cKO offspring (Extended Data Fig 8a), suggesting that different USV metrics may be driven by different mechanisms. Corroborating this, in female mHFD cKO offspring, we saw significant improvement of USV similarity (Extended Data Fig 8b) but no change in the number of USVs. *Tlr4* cKO resulted in a partial rescue of total call time, inter-syllable interval, and mean syllable number in female mHFD cKO mice, but no rescue of mean call length or mean frequency (Extended Data Fig 8b). Juvenile female mHFD cKO offspring showed a complete rescue of social preference (Fig 3e, Extended Data Fig 8f), suggesting that mHFD-induced *Tlr4*-dependent inflammation in females is responsible for most behavioral outcomes, through a non-5-HT mechanism.

To understand how mHFD results in activated Tlr4 signaling in fetal tissue, we used a Tlr4-reporter cell line (HEK-Dual™ mTLR4 (NF/IL8), Invivogen) to determine if mHFD tissue lysate stimulates Tlr4 signaling. Application of lysate from mLFD and mHFD placenta and brain tissue revealed robust Tlr4 activation in response to mHFD versus mLFD tissue (Fig 3f-g). As dietary fats reach the fetus through the placenta, and some literature suggests that saturated fatty acids (SFAs) stimulate Tlr4 signaling^53^, we determined if the increased SFAs in our model could directly activate Tlr4 signaling by stimulating the reporter cell line with six SFAs that were most abundant (and most increased) in our HFD model. While stearic acid was able to significantly activate Tlr4 signaling, as previously described^54^, the magnitude of the response was minimal (Extended Data Fig 7k).

Previous literature has demonstrated that pre-gravid obesity and high-fat diet increase circulating endotoxin levels^55–57^, and that LPS injected into pregnant dams can reach fetal tissue^58^. As Tlr4 is classically activated by bacterial endotoxins, we hypothesized that mHFD offspring tissue may be exposed to and accumulate endotoxin. Thus, we quantified bacterial endotoxin presence by limulus amebocyte lysate (LAL) assay. We detected significantly more endotoxin in mHFD placenta and fetal brain tissue from male and female offspring (Fig 3h-i), suggesting that mHFD increases endotoxin load in fetal tissue and activates Tlr4 signaling.

### Maternal overnutrition causes decreased prenatal 5-HT in human male brain tissue

To determine to what degree our findings are translatable to the human population, we obtained a large (37 individuals) cohort of matched tissues (fetal brain, fetal placenta, and maternal placenta (decidua)) from fetuses collected via elective termination at 72-82 days post conception (d.p.c.), a timeframe that closely matches the embryonic developmental window used in our mouse studies (14.5-16.5 d.p.c.; Fig 4a). PCR for the Y chromosome (*SRY;* Supplementary Table 2) identified 17 male and 20 female fetal tissue sets, with the average age of 78.2 d.p.c. for both sexes (Extended Data Fig 9a). No maternal data were available, so we assessed if triglyceride accumulation in the placenta could serve as a proxy for maternal dietary fat consumption, given that in humans, placental triglyceride/lipid levels have been shown to be increased in the context of maternal obesity^59–61^. To show proof of principle in our mouse model, triglyceride measurements from mLFD and mHFD placenta revealed a significant increase in triglyceride levels in mHFD offspring placenta, in agreement with previous literature^62^ (Extended Data Fig 9b), and placental triglyceride levels were significantly negatively correlated with fetal brain 5-HT in males (Extended Data Fig 9c). Thus, in the human samples, we assessed decidual triglyceride accumulation to use as a proxy for maternal dietary fat consumption. Triglyceride levels trended higher in pregnancies with female fetuses, but there was no significant difference between triglyceride accumulation in decidua associated with male or female pregnancies (Extended Data Fig 9d). Correlation of gene expression with maternal triglyceride accumulation from bulk RNA-seq from 16 matched male and 19 matched female brain and placenta samples revealed a strong dimorphic trend in gene expression. In male placenta, differentially correlated genes were predominantly over-expressed with increasing maternal triglycerides (i.e. gene expression increased as maternal triglycerides increased); notably, the opposite effect was seen in female placenta (Fig 4b). In the male brain, only 45 genes significantly correlated with maternal triglyceride accumulation (either positive or negative), whereas 781 genes were significantly correlated in female brain tissue (Fig 4b). At collection, the entire fetal brain tissue was collected and thus we could not control for region-specificity. To address this, we used the Allen Brain Atlas Developmental Human Transcriptome (https://www.brainspan.org/) to identify marker genes for major brain regions in human development (8-12 weeks post conception) and we plotted their gene expression (Extended Data Fig 9e). The majority of our samples showed consistent gene expression across all brain regions (i.e. one region was not oversampled). Gene Ontology (GO) Enrichment Analysis of genes significantly positively correlated with maternal triglyceride accumulation revealed enrichment of inflammatory signaling pathways in both male and female placenta (Fig 4c; full GO enrichment data is available in Supplementary Table 1). Correlation analyses of targeted genes reinforced these pro-inflammatory phenotypes; assessment of common macrophage, trophoblast and syncytiotrophoblast markers^63^ revealed specific gene associations predominantly within macrophages (Extended Data Fig 9f, Fig 4b). In agreement with our findings in mice, *TLR* expression was positively correlated with maternal triglyceride accumulation in males, along with several genes associated with MyD88-independent TLR4 signaling (Extended Data Fig 9f). In males, GO enrichment from placental genes that were negatively correlated with maternal triglycerides revealed a pattern of repressive cell differentiation, particularly of immune cells (e.g. RUNX1 differentiation and myeloid cell differentiation). That is, with increasing maternal triglyceride accumulation, gene expression of factors important for cell differentiation decreased (Fig 4c). In females, placental genes that were negatively correlated with maternal triglycerides were often involved in vascular growth and development, suggesting that maternal triglyceride accumulation may decrease placental vasculogenesis/angiogenesis (Fig 4c), an important finding given that changes in the placental vascular network are associated with numerous pathologies, such as preeclampsia^64, 65^. Inflammatory responses were additionally enriched in female brain (e.g. response to TGF-β), but too few genes were correlated in males to determine if this association was true independent of sex. These inflammatory transcriptomic patterns support our findings in mice that male mHFD offspring primarily accumulate endotoxin in placental tissue, whereas female mHFD offspring primarily accumulate endotoxin in brain tissue (see Fig 3h-i). Lastly, the top GO enrichment categories from female brain transcripts negatively correlated with maternal triglyceride accumulation involved neuronal development (Fig 4c), suggesting that maternal high-fat diet and accompanying fetal inflammation may impede brain development.

**Fig 4.**
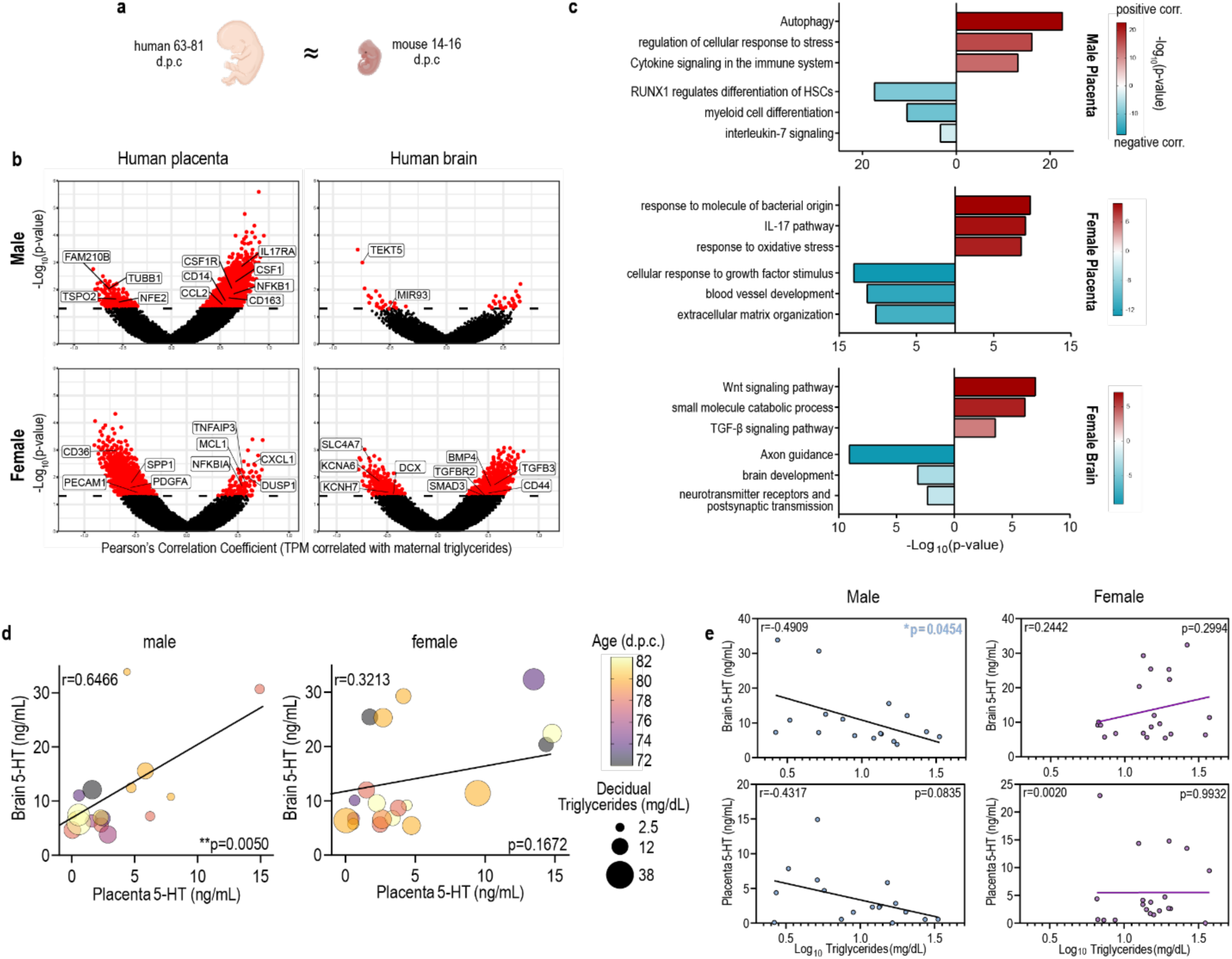
Material triglycerides negatively correlate with male fetal brain 5-HT. **a**, Human fetal development at 63-81 days post conception (d.p.c.) is roughly equivalent to mouse embryonic development at 14-16 d.p.c. **b**, Scatter plots showing transcript (TPM) correlation with maternal triglycerides versus -Log_1θ_(p-value) in male and female placenta and brain bulk sequencing data (n=16 matched male tissue sets, 19 female). Red dots indicate a significant p-value (<0.05). **c**, GO Enrichment analysis of transcripts significantly correlated with maternal triglyceride accumula­tion demonstrates robust and distinct upregulation in immune responses in male and female placenta in response to increasing maternal triglyceride accumulation. GO enrichment of transcripts significantly correlated with maternal triglyceride accumulation shows downregulation of neuronal development and function in female brain tissue in response to increasing maternal triglyceride accumulation, **d**, Multivariate dot plots show Pearson correlations between brain and placental serotonin from male and female human fetal tissues. Dot color represents the age of the matched tissue set, and dot size represents the maternal decidual triglyceride level (mg∕dL), a proxy for maternal dietary fat intake. Brain and placental serotonin levels are significantly positively correlated in male tissues between 72 and 82 d.p.c. Brain and placental serotonin levels are not correlated in female tissues (n=17 matched male tissue sets, 20 female), **e**, Brain serotonin levels are significantly negatively correlat­ed with decidual triglyceride levels in males only. Placental serotonin levels trend towards a negative correlation with decidual triglycerides in males only. Data are mean ± s.e.m.; n represents biologically independent matched human tissue set. asterisk denoted p-values are derived from Pearson’s correlations.

Finally, to determine if 5-HT levels were impacted by maternal triglyceride accumulation, we assessed 5-HT in human male and female fetal brain and placenta. Correlation analyses revealed a significant positive correlation between placental and brain 5-HT only in male fetal tissues (Fig 4d). This is consistent with previous literature (sex not reported)^30^ and our findings in male mice (see Fig 2e, Extended Data Fig 5f). Importantly, brain region marker gene expression did not correlate with 5-HT levels, and the top samples that most overrepresented midbrain gene expression profiles did not drive the correlation. Correlation analyses further revealed a negative correlation between brain or placental 5-HT levels and maternal triglyceride accumulation in male tissue only (Fig 4e). This is in agreement with our findings in mice that mHFD results in decreased brain and placental 5-HT levels in male fetal tissue (see Fig 2a-b, Extended Data Fig 4d-e). In sum, in both mice and humans, maternal high-fat diet results in the perpetuation of inflammation to the fetal brain and placenta in both sexes. In males, this inflammatory response results in aberrant phagocytosis of 5-HT neurons and long-lasting decreased levels of brain 5-HT and decreased reward behavior.

## Discussion

Here, we demonstrate a fundamental, sex-biased mechanism through which maternal high-fat diet increases offspring susceptibility to neuropsychiatric disorder development. Our results demonstrate that, in the context of maternal high-fat diet, endotoxin accumulation mediates *in utero* inflammation through the pattern recognition receptor Tlr4 in males and females, leading to increased macrophage reactivity in both the placenta and the fetal brain. In female mice, Tlr4-dependent inflammation causes diminished social preference through a 5-HT-independent mechanism. In human females, maternal decidual triglyceride accumulation is negatively associated with brain development, suggesting that Tlr4-dependent inflammation impacts neuronal development in females, though the target neuronal population is not yet known. In male mHFD offspring, embryonic microglia aberrantly phagocytose 5-HT neurons in the DRN, leading to diminished brain 5-HT from embryonic stages through adulthood, and offspring anhedonia. In human males, maternal decidual triglyceride accumulation is associated with pro-inflammatory signaling pathways in both sexes, and negatively correlated with brain 5-HT levels in males only, reinforcing our findings in mice.

Seminal work reported the placenta as a transient source of 5-HT for the fetal brain^30^, though recent studies suggest that the fetal placenta lacks the enzymes required for 5-HT synthesis^46^. No studies have investigated whether fetal sex plays a role in 5-HT synthesis or transfer in the placenta. Further, whether developmental 5-HT levels are reflective of adult 5-HT levels, and if disruptions to 5-HT levels during this window impact the brain long-term are unknown. Here, we show that fetal placenta and brain 5-HT levels are susceptible to perinatal inflammation in males only, and that decreased fetal 5-HT levels are reflective of decreased adult brain 5-HT levels in the context of maternal high-fat diet. We further found that transcript levels of both the neuronal 5-HT synthesis enzyme and 5-HT transporter (*Tph2* and *5HTT*, respectively) were decreased in male mHFD placenta, but we did not distinguish between the maternal and fetal compartments of the placenta. By manipulating fetal 5-HT levels through maternal dietary tryptophan enrichment, we can shed light on the relationship between placental and fetal brain 5-HT. While at baseline there is a significant positive correlation between placental and fetal brain 5-HT in males only, maternal tryptophan supplementation disrupted the correlation. This suggests that either the placenta is not the sole source of fetal brain 5-HT, or that the fetal brain is capable of sensing and limiting the influx of placental 5-HT in cases of excess. Alternatively, maternal dietary changes may alter the timing of the switch from placental 5-HT dependence to fetal midbrain 5-HT dependence. Careful investigation of the spatiotemporal expression of placental 5-HT transport and synthesis enzymes in male and female placenta and decidua is warranted to further clarify this topic.

5-HT has been postulated to influence macrophage phagocytic activity and cytokine production for decades^66, 67^. Here, we demonstrate a novel male-specific potential feedback loop through which microglial phagocytosis influences central 5-HT levels, which in turn influence microglial phagocytic activity. mHFD results in increased inflammation dependent on macrophage Tlr4 signaling, which results in microglial over-phagocytosis of 5-HT neurons in the DRN of males only. Considering 5-HT is thought to decrease phagocytic capacity^43^, this resulting decrease in 5-HT thus propagates the over-phagocytosis of the mHFD microglia. Both abrogation of overactive Tlr4-signaling in macrophages and increased 5-HT bioavailability via maternal tryptophan supplementation were sufficient to rescue central 5-HT levels, demonstrating the reciprocal relationship between inflammation and 5-HT in the male fetal brain.

In closing, mechanisms through which chronic maternal inflammation impact offspring neurodevelopment are scarce. In our mouse model, we demonstrate that fetal inflammation is propagated by macrophage-specific Tlr4 in both male and female mHFD offspring, along with sex-specific neurodevelopmental and behavioral outcomes. Further, we generated a large sequencing dataset of human fetal tissue from both male and female pregnancies, which demonstrated 1) a high degree of fetal inflammation in both male and female placenta, and particularly in female brain in response to increasing maternal triglyceride accumulation, and 2) sex-specific inflammatory responses. These data will shed light on which other fetal inflammatory pathways might be responsive to maternal triglyceride accumulation and will provide direction for future studies.

## Methods

### Animals

All procedures relating to animal care and treatment conformed to Duke Institutional Animal Care and Use Committee (IACUC) and NIH guidelines. Animals were group housed in a standard 12:12 light-dark cycle. The following mouse lines were used in this study: *C57BL/6J* (Jackson Laboratory, stock no. 000664), *C57BL/6NCrl* (Charles River, #027), Cx3cr1-Cre BAC-transgenic^51^, *Tlr4^f/f^* (JAX 024872^52^). Cx3cr1-Cre mice were developed in *C57BL/6N* mice but have been backcrossed with *C57BL/6J* mice for multiple generations. The Cre transgene was maintained in males for all experimental studies. Jackson Labs maintains the *Tlr4^f/f^* mice on a *C57BL/6J* background, though single nucleotide polymorphism (SNP) analysis suggests that the original mice provided to JAX were a mixed *C57BL/6N;C57BL/6J* strain. Our in-house breeding has backcrossed them with *C57BL/6J* for multiple generations.

### Diets

Female *C57BL/6J* mice were placed on randomly assigned diet at 4-weeks old and given *ad libitum* access to their assigned diet chow and water. Weight gain was assessed three times/week. After 6-weeks on diet, females were mated with *C57BL/6J* males. Pregnancy was determined by the presence of a copulation plug (gestational day 0.5) and maternal weight was assessed daily. Dams were maintained on their assigned diet chow throughout gestation and until the pups were weaned. At weaning, offspring were group housed with sex-matched littermates and provided *ad libitum* access to rodent chow 5001 (standard rodent chow, 13% kcal from fat). Low-fat, high-fat, low-fat+tryptophan, and high-fat+tryptophan diets were purchased from Research Diets (low fat: D12450Hi, high-fat: D12451i). Tryptophan enriched diets were custom formulations resulting in 1% final tryptophan content.

### Tissue Collection and Analyses

#### Murine Tissue collection

Pregnant dams were euthanized using CO_2_ following Duke University Animal Care and Use guidelines. Uterine horns were dissected and placed on ice in sterile 1X PBS. Individual embryos were separated, and placenta, brain, and tail tissue (for genotyping) were collected. Placenta were dissected into three equal pieces. One piece was flash-frozen in liquid nitrogen at stored at -80°C until ELISA. One piece was placed into 4% paraformaldehyde in PBS (PFA, Sigma) for immunohistochemistry (IHC). The final piece was placed in TriZol and stored at -80°C. Dissected fetal brains were cut in half coronally; fetal forebrain tissue was flash frozen for ELISA while the remainder of the brain was placed into 4% PFA for IHC. For adult brain collections, offspring were anesthetized with CO_2_ and transcardially perfused with saline. Further dissections were done to isolate the midbrain, and brain tissues were immediately flash frozen in liquid nitrogen and stored at -80°C until processing.

#### Microglia isolation

Microglia were isolated using CD11b-beads as previously described^68^.

#### Human tissue collection

Human placenta (maternal decidua separated from fetal placenta) and fetal brain tissues were obtained from the NIH-supported (5R24HD000836) Laboratory for the Study of Embryology at the University of Washington, Seattle between the years 1999 and 2010 and used under a protocol approved by the Duke University Institutional Review Board (Pro00014066). Fetuses with known chromosomal abnormalities were not collected. At the time of collection, tissue specimens were placed in cryogenic tubes and immediately frozen on dry ice and stored at -80°C until required for RNA or protein extraction. Prior to collection, informed consent was obtained from all individuals undergoing the elective termination.

#### Immunohistochemistry

Placenta and brain tissue were fixed in 4% PFA overnight at 4°C, cryoprotected in 30% sucrose + 0.1% sodium azide in PBS (Sigma), and embedded in OCT (Sakura Finetek, Torrance, CA) before being cryo-sectioned directly onto Superfrost slides (Fisher) at 40µm. Sections were frozen at -20°C for storage. For IHC, sections were permeabilized in 1% Triton X-100 in PBS for 12 minutes, blocked at room temperature (RT) with 5% goat serum (GS) in PBS + 0.1% Tween-20, then incubated for 2 nights at 4°C with chicken anti-Iba1 (Synaptic Systems, 234 006) or goat anti-Iba1 (Novus, NB100-1028), rat anti-CD68 (Biolegend, 137002), and rabbit anti-5-HT (Sigma, S5545) or rat anti-F4/80 (Abcam, ab6640). Sections were then washed with PBS and incubated in the following secondary antibodies: anti-rabbit Alexa-488, anti-chicken Alexa-647 or anti-goat Alexa-647, and anti-rat Alexa-568 (brain), or anti-rat Alexa-568 (placenta; 1:200; ThermoFisher), and DAPI (100µg/mL). Ten z-stacks of 0.5µm optical thickness were taken at 20X using a Zeiss AxioImager.M2 (with ApoTome.2) from at least 5 sections (each 400µm apart) from the fetal compartment of each placenta for F4/80+ cell quantification. Ten z-stacks of 0.5µm optical thickness were also taken at 20X from at least 3 fetal brain sections/animal (each section being 200µm apart) for Iba1 and CD68 immunoreactivity. For IMARIS reconstructions, z-stacks of 0.25µm optical thickness were taken to capture entire microglia. Surface reconstructions of Iba1, CD68, 5-HT, and DAPI were done for at least 19 microglia/diet from 6 mLFD, 5mLFD+Trp, 5 mHFD, and 6 mHFD+Trp male offspring and at least 12 microglia/diet from 3 mLFD, 5 mLFD+Trp, 6 mHFD, and 5 mHFD+Trp female offspring (3 litters/diet). Reconstructions from were done for at least 16 microglia/diet from 6 mLFD control, 6 mLFD cKO, 5 mHFD control, and 3 mHFD cKO male offspring and 7 mLFD control, 5 mLFD cKO, 5 mHFD control, and 4 mHFD cKO female offspring from 7 mLFD and 5 mHFD litters. Slides with cryosectioned tissue were de-identified and the individual performing the IHC, imaging, and image analyses was blind to sex, genotype, and maternal diet at all times. Statistics were run on animal average values, not individual microglia values.

#### 5-HT ELISA

Mouse placenta and brain tissue were thawed on ice and homogenized in freshly prepared lysis buffer (final concentrations: 1mg/mL ascorbic acid, 0.147% NP-40, 1X HALT Protease Inhibitor (Thermo 78443) in 1X PBS). Samples were then centrifuged at 13,000 xg for 20 minutes; supernatant was diluted 1:2 for placenta, or undiluted for fetal brain preparations, and loaded and processed according to manufacturer instructions (Enzo, ADI-900-175). Dam blood was collected in heparinized capillary tubes and immediately transferred to Eppendorf tubes on ice before being centrifuged at 1.6 x 1000 rpm for 20 minutes at 4°C. Plasma was stored at -80°C for a maximum of 2 weeks before assessment for 5-HT according to manufacturer’s instructions (1:16 dilution).

#### Quinolinic Acid/Kynurenic Acid ELISA

Mouse placenta and brain tissue were thawed on ice and homogenized with Dounce homogenizers in 500µL 1X PBS with 1X HALT Protease Inhibitor. Samples were then centrifuged at 13,000 xg for 10 minutes and supernatant processed according to manufacturer instructions (ImmuSmol IS I-0200, IS I-0100).

#### Decidual Triglycerides

Human decidua samples (stored at -80°C) were kept on dry ice to remove ∼150mg tissue which was then thawed on ice, and homogenized in 1mL NP40 Substitute Assay Reagent (Cayman Chemical, 10010303). Homogenized samples were centrifuged at 10,000 xg for 10 minutes at 4°C and the supernatant was processed (undiluted) according to manufacturer instructions at room temperature (Cayman Chemical, 10010303). Mouse placenta samples (stored at -80°C) were thawed on ice and ∼40mg of tissue was homogenized in 500µL NP40 Substitute Assay Reagent. Homogenized samples were centrifuged at 10,000 xg for 10 minutes at 4°C and the supernatant was processed (undiluted) according to manufacturer instructions at room temperature (Cayman Chemical, 10010303).

#### High-performance liquid chromatography (HPLC) with electrochemical detection

Female *C57BL/6NCrl* (Charles River #027) mice were placed on LFD or HFD as described above, with the exception that these diets were not irradiated. Pregnant females were euthanized at gd14.5 by lethal i.p. injection with ketamine/xylazine. Placentae were separated from each fetus, the amniotic sac and maternal membranes were removed, and the resulting discoid placenta was cut in half sagitally. One half was snap-frozen in liquid nitrogen for HPLC analysis. Fetal brains were dissected and snap-frozen in liquid nitrogen as well. Placenta and brain samples were analyzed for 5-HT, the 5-HT precursor tryptophan (Trp), and the 5-HT metabolite 5-hydroxyindoleacetic acid (5-HIAA). On the day of analysis, 500µl of ice-cold standard buffer (0.5 mM sodium metabisulfite, 0.2 N perchloric acid, and 0.5 mM EDTA) was added to thawed (on ice) placenta tissue and 250µl was added to thawed brain tissue. Tissues were then disrupted by sonication until completely homogenized, then centrifuged at 16,000 x g for 10 min at 4°C. The supernatant was collected and filtered through a 0.45μm membrane via centrifugation (Durapore PVDF centrifugal filters, Millipore, Billerica, MA, USA). The filtrate was kept on ice until analysis. Processed samples were separated using a 100 x 4.6 mm Kinetex (C18 5 μm 100 Å, Phenomenex) column on a reverse-phase HPLC system with a BAS LC-4B electrochemical detector with dual 3mm carbon electrode (MF-1000) and reference electrode (MF-2021) as previously described^69^. An external standard curve of all analytes was run each day. Trp quantification was performed using a mobile phase consisting of 8% acetonitrile (v/v), 0.05 M citric acid, 0.05 M Na2HPO4•H2O, and 0.1 mM EDTA. No correction for pH was needed. The detector was set to 0.875 V versus Ag/AgCl reference electrode, sensitivity at 20 nA, and a flow rate of 1.0 ml/min. 5-HT and 5-HIAA were separated with a mobile phase consisting of 18% methanol (v/v), 0.1M sodium phosphate, 0.8 mM octanesulfonic acid (anhydrous), and 0.1 mM EDTA (final pH adjusted to 3.1). The detector was set to 0.70 V, sensitivity at 20 nA, and a flow rate of 1.0 ml/min.

#### Limulus amebocyte lysate (LAL) assay for endotoxin detection

Fetal placenta and brain tissue from e14.5 mLFD and mHFD offspring were collected in a sterile environment and frozen in the vapor phase above liquid nitrogen before being transferred to 2mL Eppendorf tubes and being stored at -80°C to prevent endotoxin leaching into the tubes. The day before sampling for endotoxin, the samples were rapidly transferred to glass Dounce homogenizers and homogenized in 750µL of endotoxin free water (Lonza W50-100). Homogenized samples were then diluted 1:5 in endotoxin free water in pyrogen-free glass tubes (Lonza N207) and tested for endotoxin in duplicate using a Kinetic-QCL™ LAL Assay (Lonza, 50-650U) according to manufacturer’s instructions and including a third ‘spiked’ sample from each tissue (5EU/mL endotoxin added) to ensure acceptable recovery from every sample. Plates were run on a kinetic SPECTROstar Nano (BMG Labtech) plate reader and analyzed with MARS data analysis software (BMG Labtech) with a baseline correction of ΔO.D.=0.2^70^. Samples in which the ‘spiked’ sample fell outside the acceptable recovery range were excluded (two samples in total).

### Behavior

Behaviors are presented sex-stratified as they are often sex-dependent in their presentation^71, 72^.

#### Ultrasonic Vocalizations (USVs)

At P8, pups were brought in their home cage (with the dam being the only adult in the cage) to the testing room. Each pup was removed from their home cage and placed directly in a small cup in an insulated chamber with an AviSoft Condenser ultrasound microphone (Avisoft-Bioacoustics CM16/CMPA) suspended four inches above the contained pup. The dam and any remaining littermates were removed from the testing room in the home cage. After 3 minutes of recording, pups were weighed and sexed before being returned to their home cage. After the dam attended to each returned pup, the home cage was returned to the colony room. USV analysis was done with Avisoft and MUPET^23^ (Mouse Ultrasonic Vocalization Extraction Tool). Default MUPET parameters were used with the exception of: noise-reduction = 8.8, minimum-syllable-duration = 2.0, as we have previously optimized^39^. 80-unit repertoires were generated for each dataset (e.g. all male mLFD calls from one cohort were combined to generate ranked syllable units by frequency of occurrence) and manually inspected before USV similarity was determined using MUPET. USV similarity represents an overall Pearson’s rank order comparison of individual repertoire units (RUs) across datasets based on their spectral shape^23^.

#### Open Field Test

Adult offspring were placed in a 50cm x 50cm square enclosure with 38cm high walls and allowed to explore freely for ten minutes^73^. Behaviors were recorded and analyzed using EthoVision video tracking software (Noldus). Center-avoidance was assessed by comparing time spent at the periphery to time spent centrally/number of central entries (middle third of chamber).

#### Sociability/Social Novelty Preference

Mice were assessed for social preference or social novelty preference using a classic 3-chambered social preference test^74^. One day prior to testing, subject and stimulus animals were habituated to the testing room for a minimum of 1 hour before being individually habituated to the 3-chambered social preference boxes for 5 minutes each. On test day, subject and stimulus animals were habituated to the testing room for a minimum of one hour before testing. Juvenile offspring were tested for preference to interact with either a novel age- and sex-matched conspecific or an inanimate novel object (rubber duck). Mice were placed in the middle chamber with the inanimate object confined in a clear plexiglass cylindrical cage (plexiglass rods for thorough cleaning) on one side of the test and a novel conspecific confined in an identical cage on the other side for 5 minutes. The inanimate object was then replaced with a new novel age- and sex-matched conspecific and experimental animals were allowed to freely investigate either the new animal or the familiar animal. The entire 10-minute test was recorded with a Logitech webcam, and all behavior was manually quantified using Solomon Coder by an observer blind to sex/treatment. The time spent, latency to first instance, and frequency of each of the following behaviors was coded: in chamber, investigating (nose poking into the cage), and climbing (front paws on cup and back fully extended/arched or on top of the cage). A mouse was considered “in” a chamber after its head crossed the threshold from one chamber to the next. Mice that spent >20% of the test climbing were excluded from analysis. Mice were also excluded from analysis if they did not enter any one of the three chambers, or if the latency to enter any one chamber was greater than 150s (half the test). Social preference is represented by a sociability score significantly higher than chance (50%). Sociability score formula: (time investigating social stimulus/ (total investigation time (social + object))*100). Social Novelty Preference score formula: (time investigating novel social stimulus/ (time investigating novel + familiar social stimuli))*100).

#### Sucrose Preference

Adult offspring were tested for their preference for a 1% sucrose solution in water or plain water. Mice were singly housed and provided with 2 drinking bottles and extra enrichment (toy block or bone) for a total of 6 days. After 3 days (acclimation to 2 bottles with drinking water in both), one bottle was filled with a 1% sucrose solution while the other was filled with drinking water. Sucrose and water intake were measured daily (by weight) over the next 3 days. The positions of the bottles were switched daily to reduce any bias in side-preference. Sucrose preference is calculated as ((sucrose intake)/(sucrose + water intake))*100) averaged over 3 days.

#### Forced Swim Test

Adult offspring were assessed for mobility behavior in a forced-swim test (FST). Individual mice were placed in a cylindrical container (20cm in diameter x 50cm high) filled 2/3 with water (26°C-30°C) for 7 minutes. Mice were monitored via webcam for the testing period, and mice were removed from the test early (and not used for analysis) if their head dipped fully below the water at any point. After the test period, animals were moved to an enclosure with a small towel and heating pad to dry. Time spent time immobile (not swimming or climbing) was quantified using Solomon Coder by an observer blind to sex/treatment. Velocity and distance moved were quantified by EthoVision XT tracking software (Noldus).

### qRT-PCR

RNA was isolated using TRIzol RNA extraction reagent chloroform extraction. Either 1µg (placenta) or 200-250ng (isolated microglia) RNA was reverse-transcribed into cDNA using a Qiagen QuantiTect Reverse Transcription kit (205311). qPCR was performed on an Eppendorf Realplex ep Mastercycler using Sybr/Rox amplification with a QuantiFAST PCR kit (204056). Primer sequences are listed in Supplementary Data Table 2. Fold change was calculated using the 2^(-ΔΔCT)^ method with 18S as an endogenous control. When transcripts were not detected a CT value of 40 (max cycle number) was assigned to that sample. *Trem2* was largely undetected in e14.5 microglia (∼40% of samples equally between male and female mLFD and mHFD microglia) and is thus not represented in Extended Data Fig 7b).

### Sequencing

RNA from human tissue was prepared for sequencing using an Illumina TrueSeq RNA Exome kit and sequenced on an Illumina NovaSeq 6000 configured for a S2 flow cell at 50bp PE. Brain and placenta count matrices were processed separately. Reads were aligned to the human genome using Bowtie2^75^. Transcripts per million (TPM) was calculated on the raw counts and used for all subsequent analyses. Gene inclusion was determined using EgeR^76^ (threshold of 1). Pearson’s correlation coefficients and p-values were calculated for each gene (TPM correlated with log10(maternal triglyceride accumulation)), and data was plotted in a volcano plot using custom R script. All code for processing is currently hosted and publicly available (https://github.com/bendevlin18/human-fetal-RNASeq). Gene Ontology (GO) enrichment analysis (Metascape^77^) was performed on genes with significant positive and negative correlations separately (p<0.05).

All data is publicly available (GEO Accession number: GSE188872).

### Statistics

Statistical tests, excluding RNA-sequencing analyses, were performed using Graphpad Prism 9. Raw data as well as a description of the tests and results (including multiple comparison corrections and post-hoc analyses) are provided. Unless otherwise noted, data are mean ± s.e.m except for box plots (whiskers are min. to max., hinges of boxes are 25^th^ and 75^th^ percentiles, middle line is the median). All analyses were performed by an individual blind to animal sex, condition (maternal diet), and/or genotype.

In addition to correlation analyses in human tissues (see Fig 4e), we additionally ran a global ANCOVA to address the variance between the linear regressions by sex and tissue type. ANCOVA analysis confirms that the correlation between male fetal brain 5-HT levels and maternal triglyceride accumulation is significantly different than the correlations between male placental 5-HT levels and maternal triglyceride accumulation. It is also different than the correlations between female 5-HT levels and maternal triglyceride accumulation.

## Acknowledgements

We thank the Duke University School of Medicine for the use of the Sequencing and Genomic Technologies Shared Resource, which provided library preparation and sequencing service. We also thank Kristina Sakers for guidance and R scripts for sequencing analysis.

## Author Contributions

S.D.B, L.S., and J.B. conceived the study and, together with A.M.C, designed the experiments. A.M.C, B.A.D, J.B., L.A.G., YC.J, C.H., B.P., K.W., C.L.S., F.J., A.B.C-S, and E.R.L. performed experiments and data analysis. A.M.C. wrote the manuscript with contributions from all of the authors.

The authors declare no competing interests.

**Supplementary Information** is available for this paper.

**Supplementary Table 1**. Raw GO enrichment analysis results

**Figure1-4_RawData_Stats.** Raw data and detailed statistical information for Figures 1 through 4

**Extended_Data_Fig1-9_RawData_Stats.** Raw data and detailed statistical information for Extended Data Figures 1 through 9

**Supplementary Table 2.** PCR Primer sequences

