## Extended Data Figures for "Maternal diet disrupts the placenta-brain axis in a sex-specific manner"

**Extended Data Fig. 1 Maternal high-fat diet increases maternal and offspring weight, but does not impact litter size, sex ratio, or maternal care**

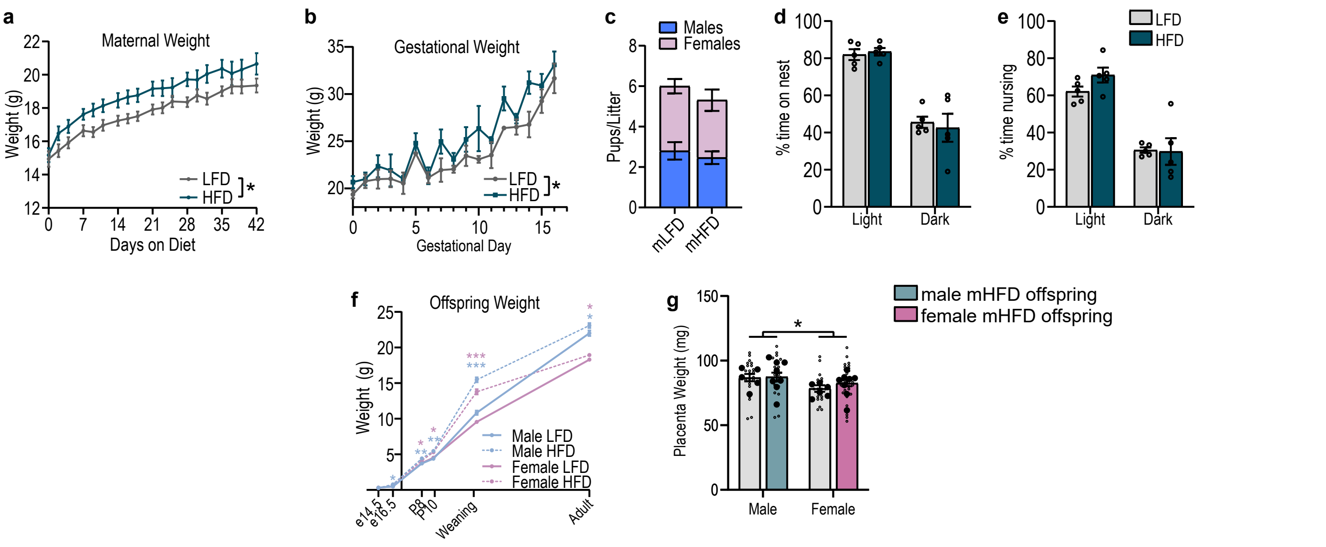


**a,** mHFD increases weight before mating (n=20 females/diet) **b,** mHFD dams are heavier during gestation than mLFD dams (n=15 dams/diet). **c,** Litter size and sex ratio are not impacted by mHFD (n=15 mLFD , 13 mHFD litters). **d,** Percent of time spent on nest is unchanged by HFD (n=5 dams/diet) **e,** Percent of time spent nursing is unchanged by HFD (n=5 dams/diet). **f,** Male and female mHFD offspring weights are increased compared to sex-matched mLFD offspring (n values can be found in ExtendedDataFig1_RawData_Stats). **g,** Placenta weight is not changed by mHFD (n=6mLFD and 8mHFD litters (large solid circles); individual placenta weights are represented in small open circles)). Data are mean ± s.e.m. asterisk denoted p-values are derived from mixed-effects two-way ANOVA (diet x time) (**a, b**), two-way ANOVA (time of day x diet) (**d, e**) (sex x maternal diet) (**g**), or unpaired two-tailed t-tests (**c, f**).

**Extended Data Fig. 2 Maternal high-fat diet imparts sex-specific offspring behavioral outcomes**

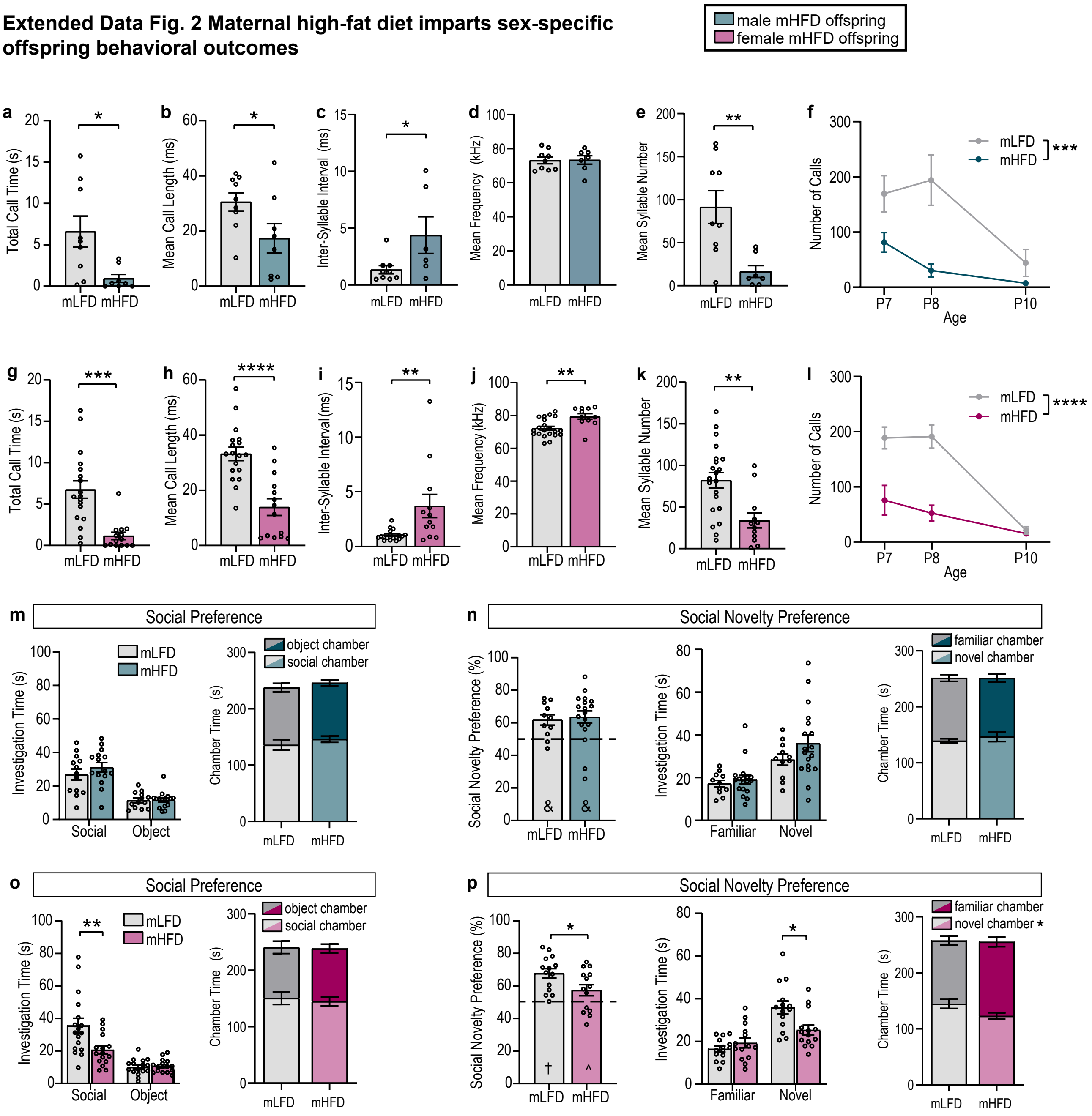


**a-e,** mHFD decreases total USV call time and mean call length, increases mean inter-syllable interval, does not affect mean call frequency, and decreases mean syllable number in male offspring (n=9 mLFD, 8 mHFD male offspring from 4 mLFD and 3 mHFD litters). **f,** mHFD dependent decrease in USV number is apparent throughout neonatal development in male offspring (P8 data as previously shown, n= 11 P7 and 7 P10 mLFD offspring from 3 P7 litters and 4 P10 litters, 14 P7 and 8 P10 mHFD offspring from 4 P7 and 3 P10 litters). **g-k,** mHFD decreases total USV call time and mean USV call length, increases mean inter-syllable interval and mean call frequency, and decreases mean syllable number in female offspring (n=18 mLFD, 14 mHFD female offspring from 5 mLFD and 3 mHFD litters). **l,** mHFD dependent decrease in USV number is apparent throughout neonatal development in female offspring (P8 data as shown previously, n=13 P7 and 14 P10 mLFD offspring from 5 P7 and 5 P10 litters, 5 P7 and 10 P10 mHFD offspring from 3 P7 and 3 P10 litters) **m,** mHFD does not alter male offspring social or object investigation times or chamber times during a 3-chamber social preference test (n=13 mLFD, 15 mHFD male offspring from 5 mLFD and 6 mHFD litters). **n,** Male mLFD and mHFD offspring display a strong preference for a novel social stimulus over a familiar one. mHFD does not alter male offspring novel or familiar investigation times or chamber times in a social novelty preference test (n=11 mLFD and 18 mHFD offspring from 4 mLFD and 7 mHFD litters). **o,** Female mHFD offspring spend less time investigating a social stimulus than female mLFD offspring, but overall chamber time is not different (n=17 mLFD, 16 mHFD female offspring from 6 mLFD and 6 mHFD litters). **p,** Female mHFD offspring have decreased preference for a novel social stimulus compared to mLFD female offspring. Female mHFD offspring spend less time investigating a novel social stimulus than female mLFD offspring, and spend less time in the chamber containing the novel conspecific (n=14 mLFD and 14 mHFD offspring from 5 mLFD and 5 mHFD litters). Data are mean ± s.e.m.; asterisk denoted p-values are derived from unpaired two-tailed t-tests (**a, b, c, e, g, h, i, j, k, o, p**) or 2-way ANOVA (diet x age; **f, l**). other p-values († $p<0.001$, & $p<0.01$, ^$p<0.05$) are derived from one-sample t-tests assessing difference from chance (50%) (**n, p**).

**Extended Data Fig. 3 Maternal high-fat diet imparts sex-specific offspring behavioral outcomes**

male mHFD offspring
female mHFD offspring

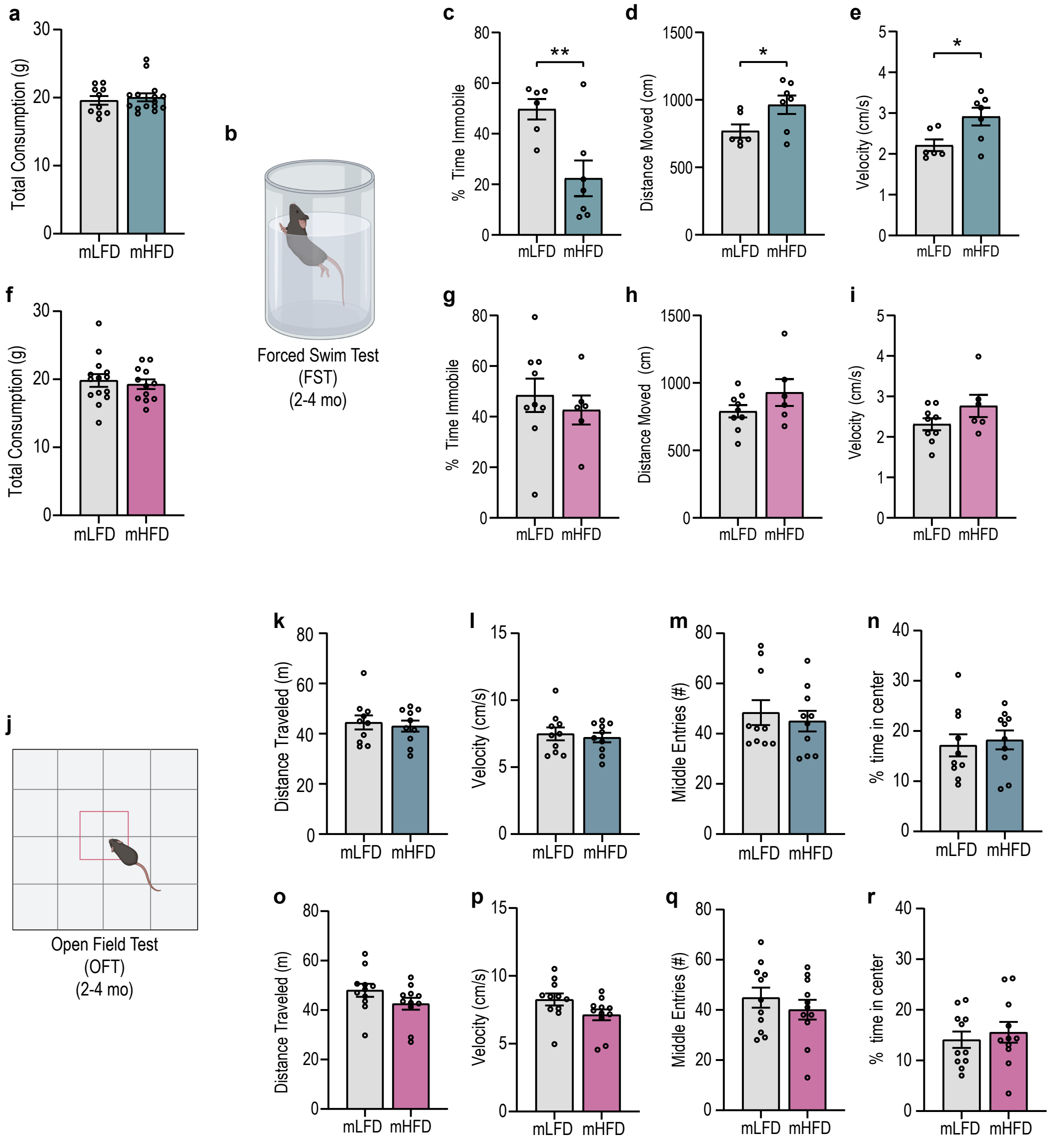


**a,** Male offspring total consumption (water + sucrose) is unaffected by maternal diet (n=10 mLFD, 15 mHFD male offspring from 5 mLFD and 5 mHFD litters). **b,** Schematic of forced swim test (FST). **c-e,** Male mHFD offspring spend less time immobile than mLFD offspring, move a greater cumulative distance, and have a higher swimming velocity than mLFD offspring during an FST (n= 6 mLFD and 7 mHFD offspring from 3 mLFD and 4 mHFD litters). **f,** Female offspring total consumption is unaffected by maternal diet (n=14 mLFD, 12 mHFD female offspring from 5 mLFD and 5 mHFD litters). **g-i,** Female mHFD perform similarly to female mLFD offspring in a forced swim test (n=9 mLFD, 6 mHFD offspring from 4 mLFD and 3 mHFD litters). **j,** Schematic of an open field test. **k-n,** Distance traveled, velocity, number of middle entries, and percent of time in center in an open field test are unchanged in male mHFD offspring (n=10 mLFD, 10 mHFD male offspring from 3 mLFD and 4 mHFD litters). **o-r,** Distance traveled, velocity, number of middle entries, and percent of time in center in an open field test are unchanged in female mHFD offspring (n=11 mLFD, 11 mHFD female offspring from 3 mLFD and 3 mHFD litters). Data are mean ± s.e.m.; asterisk denoted p-values are derived from unpaired two-tailed t-tests (**c, d, e**).

**Extended Data Fig. 4 Maternal high-fat diet decreases serotonin in male offspring**

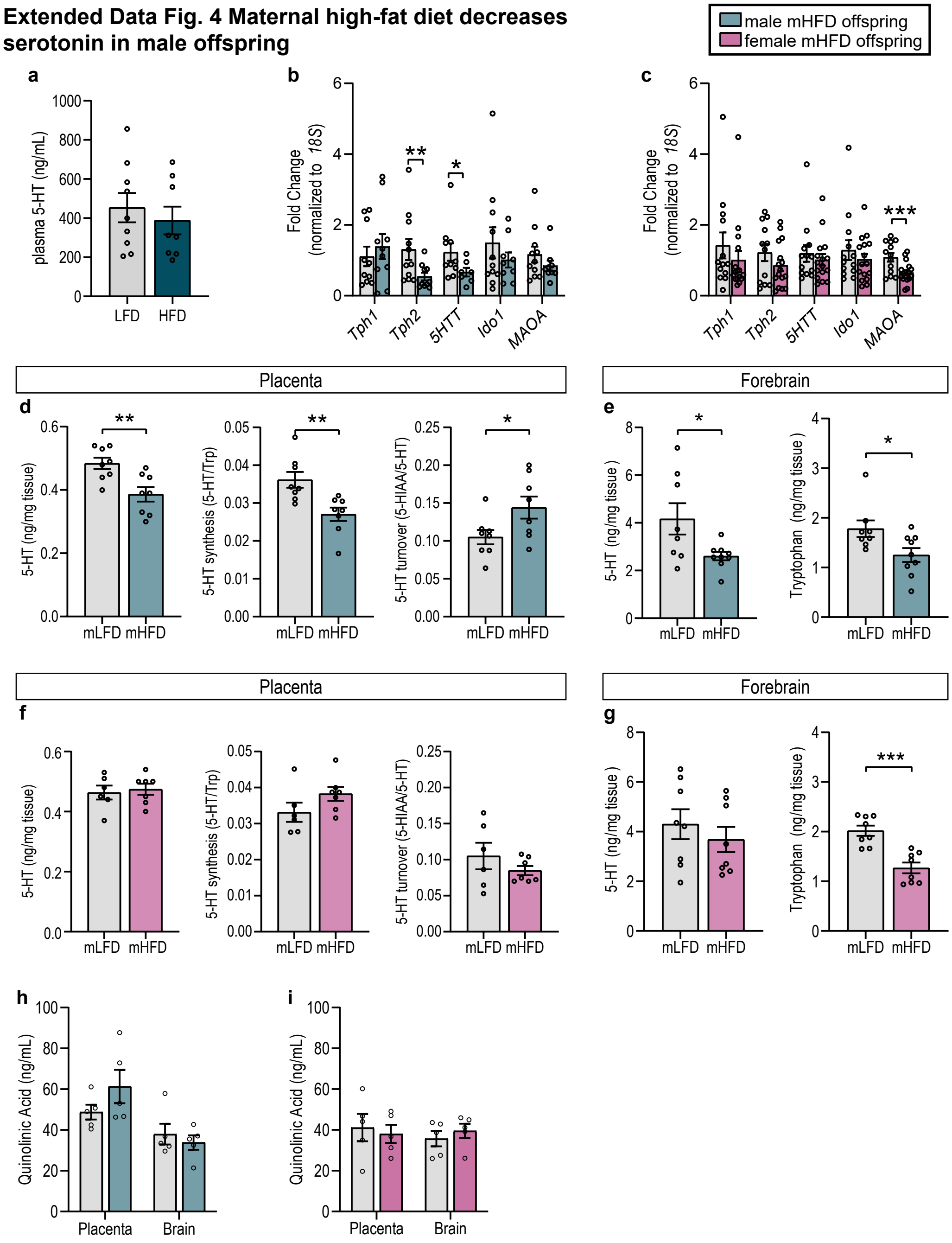


**a,** Maternal plasma 5-HT levels are unaffected by HFD at gd14.5 (n=9 LFD and 8 HFD pregnant dams). **b,** *Tph2* and *5HTT* are signficantly decreased in male mHFD placenta (n=10 mLFD, 9 mHFD offspring from 3 mLFD and 4 mHFD litters). **c,** *MAOA* is significantly decreased in female mHFD placenta (n= 13 mLFD, 16 mHFD offspring from 3 mLFD and 4 mHFD litters). **d,** 5-HT and 5-HT synthesis (5-HT/tryptophan (Trp)) are significantly decreased while 5-HT turnover (5-HIAA/5-HT) is increased in male mHFD placenta (n= 8 mLFD and 8 mHFD offspring from 8 mLFD and 8 mHFD litters). **e,** 5-HT and Tryptophan are significantly decreased in male mHFD forebrain tissue (n=8 mLFD and 9 mHFD offspring from 8 mLFD and 9 mHFD dams). **f,** mHFD does not influence 5-HT, 5-HT synthesis, or 5-HT turnover in female placenta (n=6 mLFD and 7 mHFD from 6 mLFD and 7 mHFD litters). **g,** mHFD does not influence 5-HT but decreases tryptophan levels in female forebrain (n=8 mLFD and 6 mHFD from 8 mLFD and 6 mHFD litters). **h**, Quinolinic acid levels are unaffected by mHFD in male placenta and fetal brain (n=5 mLFD and 5mHFD offspring from 5 mLFD and 5 mHFD litters). **i**, Quinolinic acid levels are unaffected by mHFD in female placenta and fetal brain (n=5 mLFD and 5 mHFD offspring from 5 mLFD and 5 mHFD litters). asterisk denoted p-values are derived from unpaired one-sample t-tests (**b, c**), or unpaired two-tailed t-tests (**d, e, g**)

**Extended Data Fig. 5 Maternal tryptophan supplementation rescues mHFD-induced behavioral changes in male offspring**

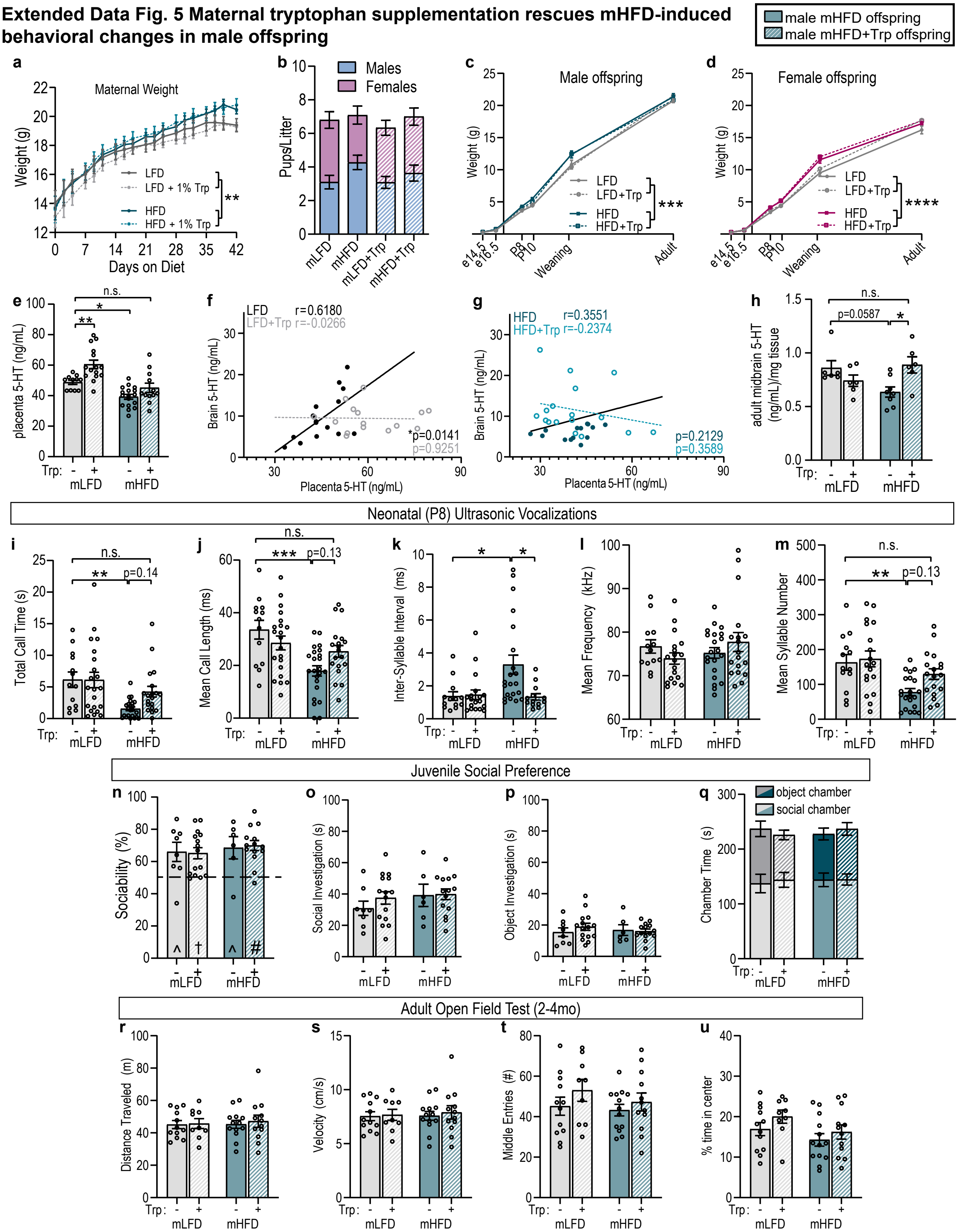


**a,** Dietary tryptophan enrichment does not impact maternal weight (n=15 LFD, 16 LFD+Trp, 15 HFD, 17 HFD+Trp females) **b,** Dietary tryptophan enrichment does not impact litter size or sex ratio (n=10 mLFD, 12 mLFD+Trp, 11 mHFD, 11 mHFD+Trp litters) **c-d,** Maternal dietary tryptophan enrichment does not impact offspring weight (n values can be found in ExtendedDataFig5_Raw_Stats) **e,** Maternal dietary tryptophan enrichment increases serotonin in mLFD, but not mHFD, male placenta (n=11 mLFD, 15 mLFD+Trp, 17 mHFD, 12 mHFD+Trp offspring from 4 mLFD, 5 mLFD+Trp, 4 mHFD, and 4 mHFD+Trp litters). **f,** Maternal dietary tryptophan enrichment disrupts the positive correlation between brain and placenta serotonin in male offspring (n=15 mLFD +/- Trp offspring from 4 mLFD and 5 mLFD+Trp litters). **g,** Brain and placenta serotonin levels are not correlated in male mHFD +/- tryptophan offspring (n=14 mHFD, 17 mHFD+Trp offspring from 4 mHFD and 5 mHFD+Trp litters). **h,** Maternal dietary tryptophan enrichment increases serotonin levels in mHFD adult male midbrain (n=6 mLFD, 6 mLFD+Trp, 8 mHFD, 6 mHFD+Trp offspring from 4 mLFD, 6 mLFD+Trp, 6 mHFD, and 6 mHFD+Trp litters). **i-m,** Total call time, mean call length, mean inter-syllable interval, mean frequency, and mean syllable number in male mLFD and mHFD +/- tryptophan offspring (n=13 mLFD, 21 mLFD+Trp, 23 mHFD, 19 mHFD+Trp offspring from 5 mLFD, 7 mLFD+Trp, 8 mHFD, and 6 mHFD+Trp litters). **n-q,** Maternal dietary tryptophan enrichment does not impact male offspring juvenile social behavior (n=8 mLFD, 15 mLFD+Trp, 6 mHFD, and 14 mHFD+Trp male offspring from 3 mLFD, 6 mLFD+Trp, 3 mHFD, and 4 mHFD+Trp litters). **r-u,** Maternal dietary tryptophan enrichment does not impact open field behavior in adult male offspring (12 mLFD, 9 mLFD+Trp, 13 mHFD, 12 mHFD+Trp male offspring from 4 mLFD, 4 mLFD+Trp, 3 mHFD, and 5 mHFD+Trp litters). Data are mean ± s.e.m.; asterisk denoted p-values are derived from 2-way ANOVA (Fat x Trp content, **b, e, h, i, j, k, m**), 3-way ANOVA (Fat Content x Trp content x Days on Diet/Age, **a, c, d**), or Pearson's correlation (**f, g**). other p-values (# $p<0.0001$, † $p<0.001$, ^ $p<0.05$) are derived from one-sample t-tests assessing difference from chance (50%) (**n**).

**Extended Data Fig. 6 Maternal tryptophan supplementation does not rescue behavior in female mHFD offspring**

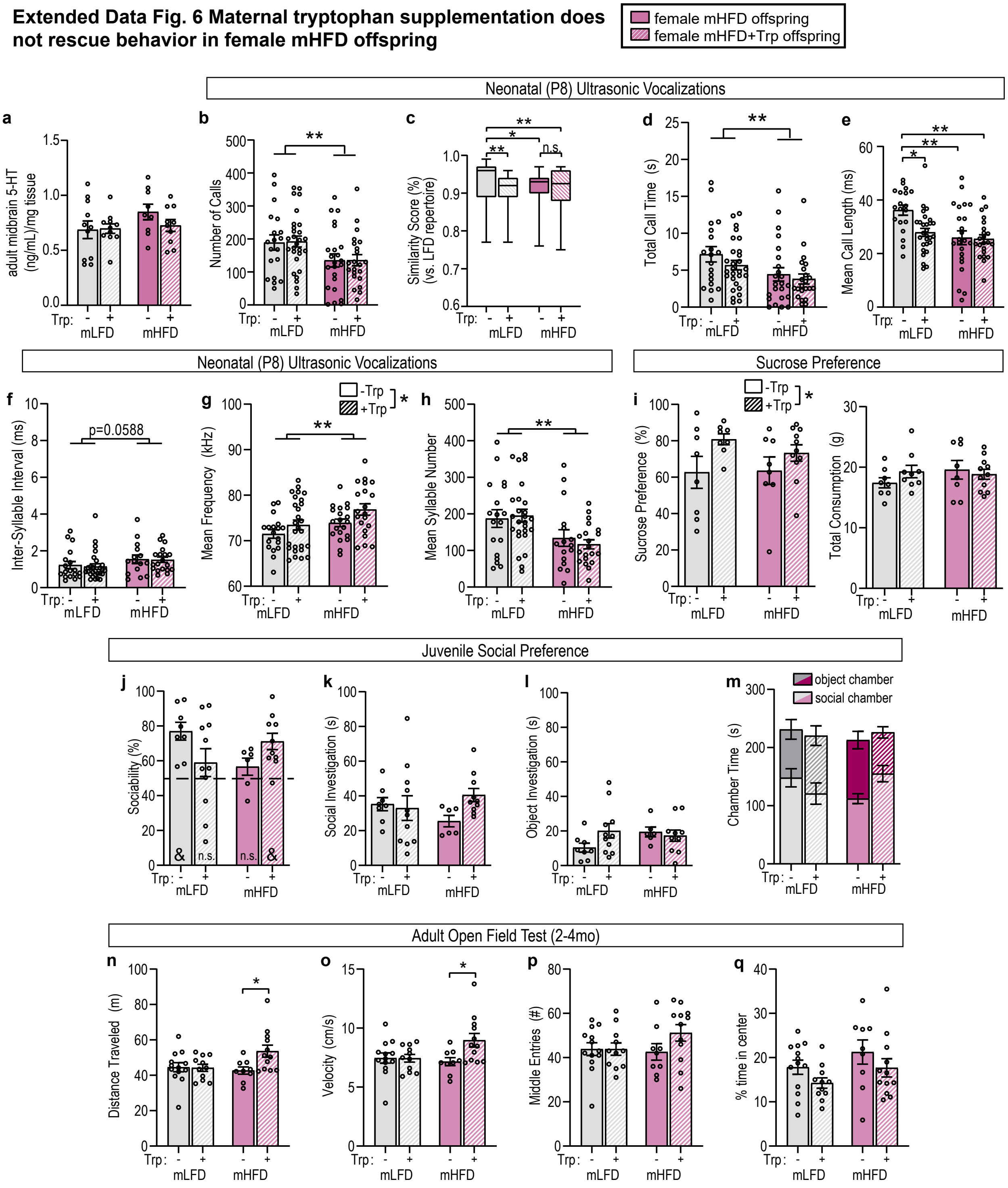


**a,** Maternal dietary tryptophan enrichment does not impact adult female offspring midbrain serotonin levels (n=11 mLFD, 11 mLFD+Trp, 9 mHFD, 10 mHFD+Trp offspring from 6 mLFD, 9 mLFD+Trp, 7 mHFD, and 8 mHFD+Trp litters). **b-h,** Maternal dietary tryptophan enrichment does not rescue mHFD-dependent changes to neonatal female ultrasonic vocalizations (n=19 mLFD, 28 mLFD+Trp, 23 mHFD, 22 mHFD+Trp offspring from 6 mLFD, 8 mLFD+Trp, 8 mHFD, and 9 mHFD+Trp litters). i, Maternal tryptophan enrichment increases female offspring sucrose preference regardless of maternal dietary fat intake (n=8 mLFD, 8 mLFD+Trp, 8 mHFD, 11 mHFD+Trp from 3 mLFD, 5 mLFD+Trp, 4 mHFD, and 6 mHFD+Trp litters) **j-m,** Maternal tryptophan enrichment infleunce on juvenile social preference (n=8 mLFD, 11 mLFD+Trp, 6 mHFD, and 10 mHFD+Trp offspring from 3 mLFD, 5 mLFD+Trp, 4 mHFD, and 3 mHFD+Trp litters). **n-p,** Maternal tryptophan enrichement influence on open field behavior (n=13 mLFD, 11 mLFD+Trp, 9 mHFD, and 12 mHFD+Trp offspring from 4 mLFD, 4 mLFD+Trp, 4 mHFD, and 4 mHFD+Trp litters). Data are mean ± s.e.m.; asterisk denoted p-values are derived from 2-way ANOVA (fat content x tryptophan content; **b, c, d, e, f, g, h, i, n, o**). Other p-values (&p<0.01) are derived from one-sample t-tests assessing difference from chance (50%) (**j**).

**Extended Data Fig. 7 mHFD induces Tlr4-dependent embryonic inflammation driving behavior changes in males and females**

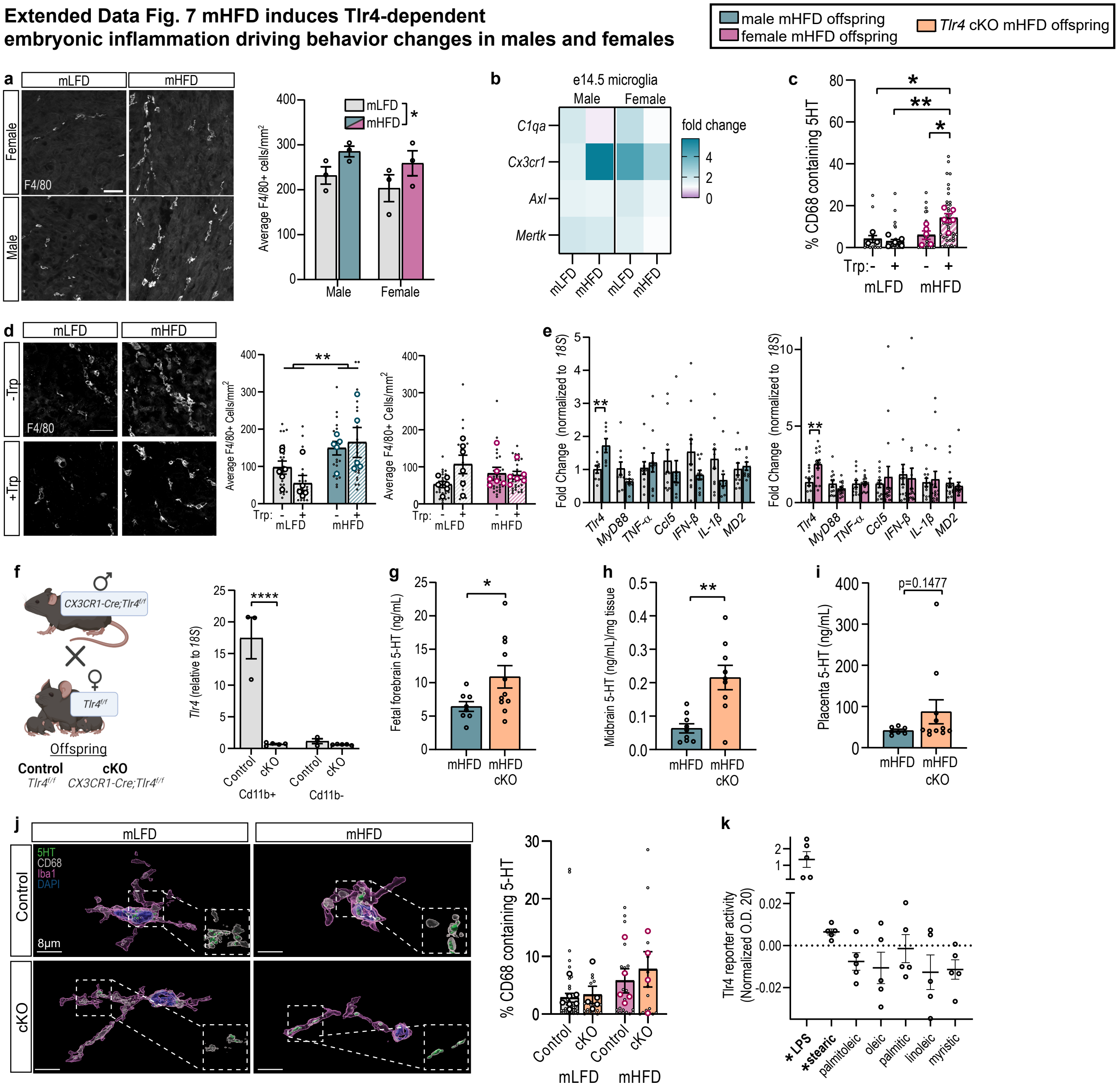


**a,** Increased macropahge density in male and female mHFD placenta (scale=50µm; n=3 animals per sex/diet). **b,** qPCR (Mean ΔΔCt values shown normalized to *18S*) from male and female e14.5 midbrain microglia (n=4 male and 5 female/diet from 4 mLFD and 5 mHFD litters) **c,** IMARIS reconstruction quantification from female e14.5 DRN microglia. Statistics shown for animal averages (large circles), small circles are individual microglia. (n=3 mLFD, 5, mLFD+Trp, 6 mHFD, 5 mHFD+Trp from 3 litters/diet). **d,** Representative images of male placenta macrophages. Macrophage density in the placenta remains increased in mHFD+Trp male offspring. Statistics shown for animal averages (large circles), small circles are individual images. (scale=50µm; n=6 mLFD, 5 mLFD+Trp, 5 mHFD, 5 mHFD+Trp). **e,** qPCR from male and female e14.5 placenta (male: n=11 mLFD, 10 mHFD; female: n=13 mLFD, 15 mHFD from 3 mLFD and 4 mHFD litters). **f,** Schematic of macrophage-specific Tlr4 knockout and confirmation of Tlr4 knockdown in microglia (n=3 control, 5 Tlr4 cKO mice). **g-h,** Serotonin levels are significantly increased by macrophage-specific loss of Tlr4 in male mHFD offspring fetal forebrain and adult midbrain (n=8 mHFD control, 11 mHFD cKO fetal forebrain from 6 mHFD litters; n=9 mHFD, 9 mHFD cKO adult midbrain from 4 litters). **i,** Placenta 5-HT in mHFD male offspring with or without macrophage Tlr4-signaling (n=6 mHFD control, 11 mHFD cKO from 6 mHFD litters) **j,** IMARIS reconstructions from female mLFD and mHFD control and cKO e14.5 DRN. Statistics ran for animal averages (large circles), small circles are individual microglia. (n=7 mLFD control, 5 mLFD cKO, 5 mHFD control, and 4 mHFD cKO from 7 mLFD and 5 mHFD litters) **k,**Tlr4-reporter activity in response to LPS and saturated fatty acids. Data are mean ± s.e.m.; denoted p-values are derived from unpaired two-tailed t-tests (**e**, **g**, **h**, **i**), two-way ANOVA (**a**, **c**, **d**, **f**), or one-sample t-tests assessing difference from 0 (baseline; **k**)

**Extended Data Fig.8 mHFD induces Tlr4-dependent inflammation driving offspring behavior changes**

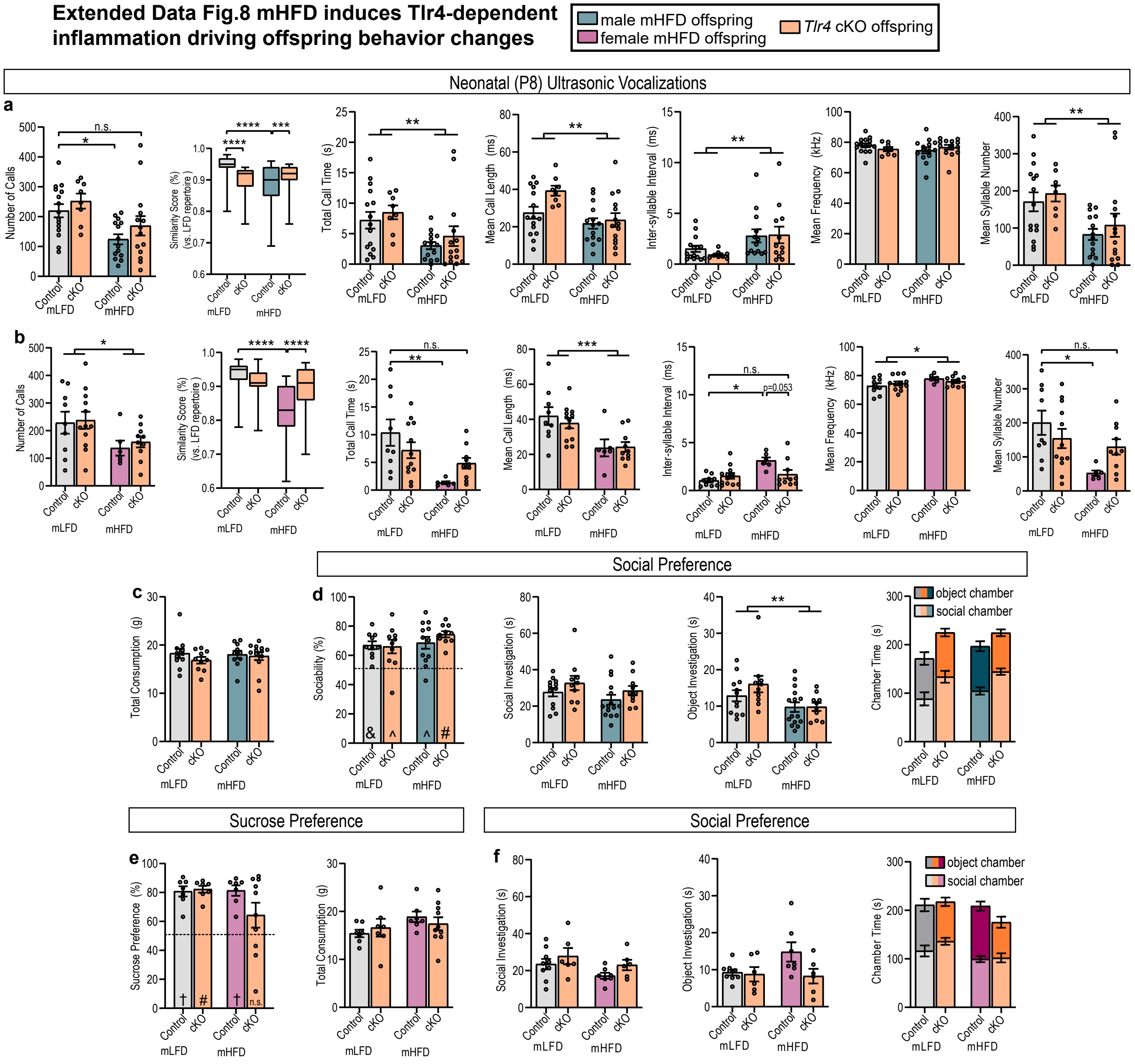


**a,** USV metrics from male and **b,** female control and cKO mLFD and mHFD offspring (n=15 male mLFD control and 8 cKO from 5 litters and 14 male mHFD control and 14 cKO from 6 litters; n=9 female mLFD control and 12 cKO from 7 litters and 6 female mHFD and 10 cKO from 5 litters). **c,** Total liquid consumption in mLFD and mHFD control and cKO males (n=11 mLFD, 10 mLFD cKO, 11 mHFD, and 10 mHFD cKO from 5 mLFD and 6 mHFD litters) **d,** Social preference in male mLFD and mHFD control and Tlr4 cKO offspring (n=10 mLFD, 10 mLFD cKO, 12 mHFD, 10 mHFD cKO from 5 mLFD and 6 mHFD litters) **e,** Sucrose preference in female mLFD and mHFD control and Tlr4 cKO offspring (n=7 mLFD, 7 mLFD cKO, 7 mHFD, 10 mHFD cKO from 5 mLFD and 4 mHFD litters) **f,** Social preference metrics in female mLFD and mHFD control and Tlr4 cKO offspring (n=9 mLFD, 6 mLFD cKO, 7 mHFD, and 6 mHFD cKO from 5 mLFD and 5 mHFD litters). Data are mean ± s.e.m.; denoted p-values are derived from two-way ANOVA (**a, b, d, f**). other p-values (# p<0.0001, † p<0.001, & p<0.01, ^p<0.05) are derived from one-sample t-tests assessing difference from chance (50%) (**d, e**).

**Extended Data Fig 9. Human maternal decidual triglyceride accumulation negatively correlates with fetal brain serotonin in males only**

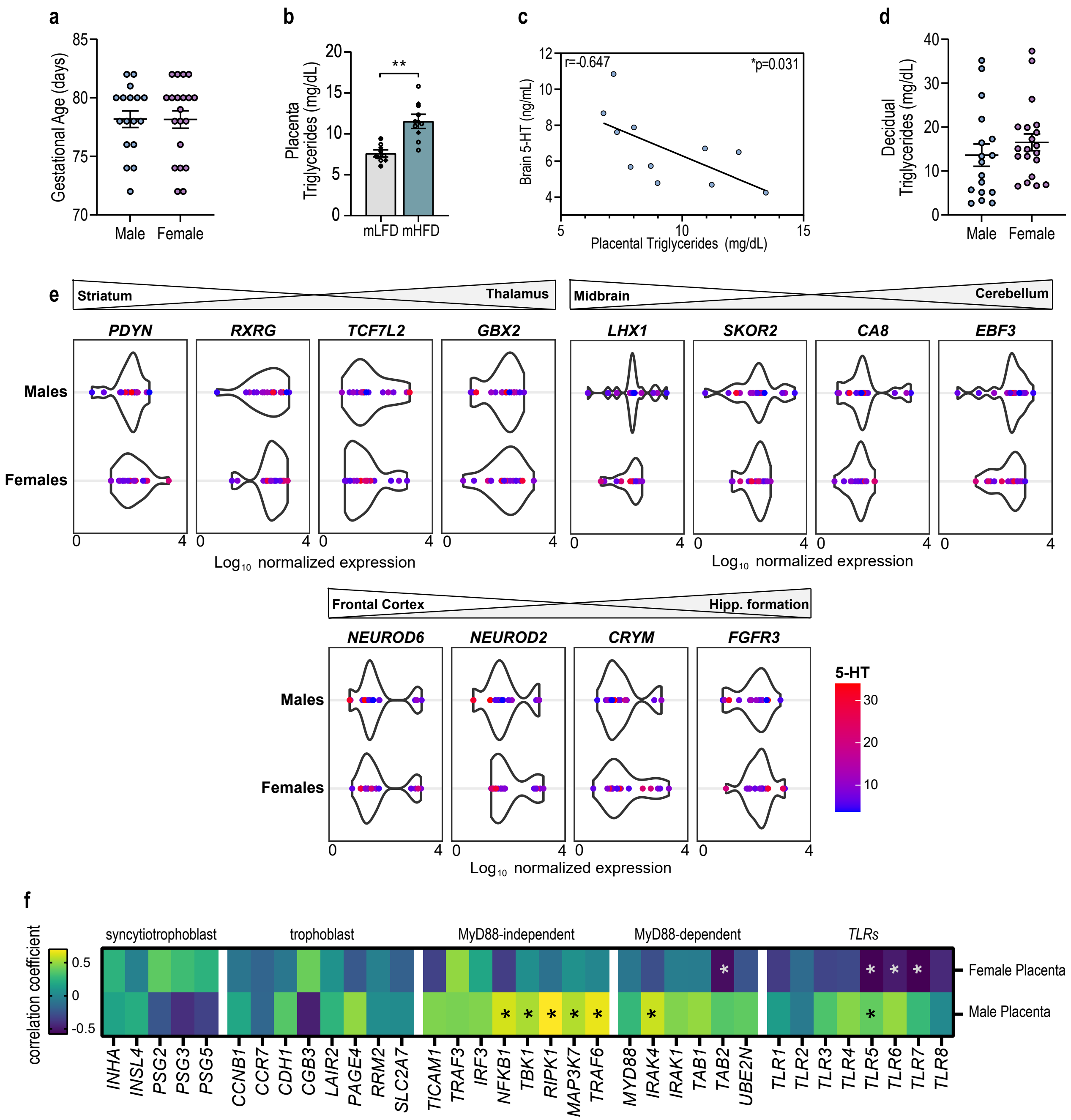


**a,** Average gestational age was equal in male and female human tissue samples. **b,** Triglyceride levels are increased in mHFD placenta (n=7 mLFD and 8 mHFD male placenta (open circles from 5 mLFD and 3 mHFD dams; statistics shown for litter average (diamonds)). **c,** Fetal brain serotonin levels are significantly negatively correlated with placental triglyceride accumulation (n=11 individuals from 5 litters). **d,** Average decidual triglyceride levels were statistically equal but trended higher in female pregnancies versus male. **e,** Violin plot showing log normalized expression of marker genes for striatum and dorsal thalamus, midbrain and cerebellum, and frontal context and hippocampal formation regions from human brain samples in males and females. The color of the dot representing each sample shows the brain 5-HT concentration for that sample. **f,** Heatmap depicting strength Pearson's correlations for select genes marking syncytiotrophoblasts, trophoblasts, and immune-related signaling in the placenta. Data are mean ± s.e.m; asterisk denoted p-values are derived from unpaired two-tailed t-tests (**b**) or Pearson's correlations (**c**, **f**).
