## Supplementary Table 2 for "Maternal diet disrupts the placenta-brain axis in a sex-specific manner"

**Extended Data Table 2. PCR Primer Sequences**

| **Target** | **Forward (5’-3’)** | **Reverse (5’-3’)** |
| --- | --- | --- |
| *Tph1* | AACAAAGACCATTCCTCCGAAAG | TGTAACAGGCTCACATGATTCTC |
| *Tph2* | CCATTGTGACCCTGAATCC | GGTGAGAGCATCTGTCTAAC |
| *5HTT* | GCTGAGATGAGGAACGAAG | GGCAAAGAATGTGGATGCTG |
| *Ido1* | GCTTTGCTCTACCACATCCAC | CAGGCGCTGTAACCTGTGT |
| *MAOA* | GCCCAGTATCACAGGCCAC | CGGGCTTCCAGAACCAAGA |
| *Tlr4* | CAGCAGAGGAGAAAGCAT | CACCAGGAATAAAGTCTCTG |
| *MyD88* | CAAGGCGATGAAGAAGGAC | CGCATCAGTCTCATCTTCCC |
| *TNF-α* | GTCGTAGCAAACCACCAA | AGAACCTGGGAGTAGACAAGG |
| *Ccl5* | GCTGCTTTGCCTACCTCTCC | TCGAGTGACAAACACGACTGC |
| *IFN-β* | CAGCTCCAAGAAAGGACGAAC | GGCAGTGTAACTCTTCTGCAT |
| *IL-1β* | GTCTTCCTAAAGTATGGGCTG | CACAGGCTCTCTTTGAAC |
| *MD2* | CGCTGCTTTCTCCCATATTGA | CCTCAGTCTTATGCAGGGTTCA |
| *Cd11b* | CTATTTGTTCGGCTCCAAC | GCATCAAAGAGAACAAGG |
| *Trem2* | CTGGAACCGTCACCATCACTC | CGAAACTCGATGACTCCTCGG |
| *C1qa* | ACCCAGGAGAGTCCATACCA | ACAGACAAAGGTCCCACTTG |
| *Cx3cr1* | TGGATTCCCAGCAAGCCATAG | GTCTGCTACCCTCACAAA |
| *Cd68* | CAACAAAACCAAGGTCCAG | CACATTGTATTCCACCGC |
| *Axl* | GGAACCCAGGGAATATCACAGG | AGTTCTAGGATCTGTCCATCTCG |
| *Mertk* | CAGGGCCTTTACCAGGGAGA | TGTGTGCTGGATGTGATCTTC |
| *18S* | GAATAATGGAATAGGACCGC | CTTTCGCTCTGGTCCGTCTT |
| *Sry* | GTGTCTCAAAGCCTGCTCTTC | CATGTACTGCTAGCAGCTATC |
| *Myog* | TTACGTCCATCGTGGACAGCAT | TGGGCTGGGTGTTAGTCTTAT |
| SRY (human) | TCAGCAAGCAGCTGGGATAC | AACTGCAATTCTTCGGCAGC |
| ATL1 (human) | CCCTGATGAAGAACTTGTATCTC | GAAATTACACACATAGGTGGCACT |
